# A novel allele of *ASY3* promotes meiotic stability in autotetraploid *Arabidopsis lyrata*

**DOI:** 10.1101/2019.12.25.888388

**Authors:** Paul J. Seear, Martin G. France, Catherine L. Gregory, Darren Heavens, Roswitha Schmickl, Levi Yant, James D. Higgins

## Abstract

In this study we performed a genotype-phenotype association analysis of meiotic stability in ten autotetraploid *Arabidopsis lyrata* and *A*. *lyrata/A. arenosa* hybrid populations collected from the Wachau region and East Austrian Forealps. The aim was to determine the effect of eight meiosis genes under extreme selection upon adaptation to whole genome duplication. Individual plants were genotyped by high-throughput sequencing of the eight meiosis genes (*ASY1, ASY3, PDS5b, PRD3, REC8, SMC3, ZYP1a/b*) implicated in synaptonemal complex formation and phenotyped by assessing meiotic metaphase I chromosome configurations. Our results reveal that meiotic stability varied greatly (20-100%) between individual tetraploid plants and was associated with segregation of a novel allele orthologous to the budding yeast RED1 chromosome axis protein, Asynapsis3 (ASY3), derived from *A. lyrata.* The adaptive ASY3 protein possesses a putative in-frame tandem duplication (TD) of a serine-rich region upstream of the coiled-coil domain that has arisen at sites of DNA microhomology. The frequency of multivalents observed in plants homozygous for the *ASY3 TD* haplotype was significantly lower than plants heterozygous for *TD/ND* (non-duplicated) *ASY3* haplotypes. Chiasma distribution was significantly altered in the stable plants compared to the unstable plants with a shift from proximal and interstitial to predominantly distal locations. The number of HEI10 foci at pachtyene that mark class I crossovers was significantly reduced in meiotic nuclei from ASY3 TD homozygous plants compared to ASY3 ND/TD heterozygotes, indicating an adaptive consequence of the ASY3 TD allele. From the ten populations, fifty-eight alleles of these 8 meiosis genes were identified, demonstrating dynamic population variability at these loci which nevertheless exhibit signatures of strong hard selective sweeps. Widespread chimerism between alleles originating from *A*. *lyrata/A. arenosa* and diploid/tetraploids indicates that this group of rapidly evolving genes provide precise adaptive control over meiotic recombination in the tetraploids, the very process that gave rise to them.

**Author summary:** Whole genome duplication can promote adaptability, but is a dramatic mutation usually resulting in meiotic catastrophe and genome instability. Here we focus on a case of coordinated stabilization of meiotic recombination in ten autotetraploid *Arabidopsis lyrata* and *A*. *lyrata/A. arenosa* hybrid populations from the Wachau region and East Austrian Forealps. We fuse population genomic data with a genotype-phenotype association study, concentrating on the effects of eight meiosis genes (*ASY1, ASY3, PDS5b, PRD3, REC8, SMC3, ZYP1a/b*) implicated in synaptonemal complex formation in the tetraploids under extreme selection. Our analysis demonstrates that a novel allele of the meiotic chromosome axis protein Asynapsis3 that contains an in-frame duplication of a serine-rich region is the major determinant of male meiotic stability. This adaptive restabilisation appears to be achieved by a reduction in the number of meiotic crossovers as well as a shift in their positioning towards the chromosome ends. Of the eight genes, fifty-eight alleles were identified, indicating dynamic population variability at these loci under extreme selection. In addition, widespread allelic chimerism between alleles originating from *A*. *lyrata/A. arenosa* and diploid/tetraploids indicates that this group of rapidly evolving genes provide precise adaptive control over meiotic recombination in the tetraploids, the very process that gave rise to them.

## Introduction

Whole genome duplication (WGD) occurs in all eukaryotic kingdoms, and is associated with adaptability, speciation and evolvability [1, 2]. At the same time, it is also one of the most dramatic mutations observed, usually resulting in catastrophic problems during meiosis, when ensuring stable chromosome segregation and genome integrity is paramount [3]. Because efficient meiosis is required for the formation of euploid gametes during sexual reproduction, selection acts strongly on standing variation from the progenitor diploids in newly arisen polyploids.

In allopolyploids (formed by both genome duplication and interspecies hybridization), loci required for correct chromosome pairing and recombination have been identified in wheat [4], oil seed rape [5, 6] and *Arabidopsis suecica* [7]. However, in autopolyploids (which form within-species, without hybridization), there has been no functional confirmation of any gene controlling correct chromosome pairing, synapsis and crossing over (CO), although we have detected clear signatures of extreme selection in eight meiosis genes associated with the synaptonemal complex (SC) (*ASY1, ASY3, PDS5b, PRD3, REC8, SMC3, ZYP1a, ZYP1b*) in the young autotetraploid *Arabidopsis arenosa* [8].

The SC is a tripartite protein structure consisting of two lateral elements and a central element, specific to meiotic prophase I that is required for normal levels of COs in the majority of sexually reproducing eukaryotes [9]. In Arabidopsis, the chromosome axes (which come to form the SC lateral elements) consist of a scaffold of cohesin proteins (SMC1, SMC3, PDS5, REC8 and SCC3)[10–14] that organise sister chromatids into a loop/base conformation [15]. PRD3, the budding yeast MER2 homolog, is required for double-strand break (DSB) formation and is not an SC protein *per se* but may juxtapose the potential DSB site with the chromosome axis to promote inter-homolog recombination [16, 17]. In *Sordaria*, MER2 also transfers and releases recombination complexes to and from the SC central region [16]. The meiosis specific proteins ASY1, ASY3 and ASY4 load onto the chromosome axis defined by the cohesin scaffold, to promote inter-homolog recombination [18–20]. ASY1 and ASY3 are the functional homologs of budding yeast HOP1 and RED1 and, HORMAD1/2 and SCP2 in mammals, respectively, that facilitate correct chromosome pairing and synapsis, required for wild-type COs [19, 21, 22]. In Arabidopsis, synapsis is initiated by installation of the transverse filament proteins ZYP1a/b between homologous chromosomes, thus ensuring appropriate levels of COs [23].

*Arabidopsis lyrata* and *A. arenosa* represent powerful models for investigating adaptation to autopolyploidy, particularly their populations from the eastern Austrian Forealps, where interspecific hybridization and introgression is frequent [8] [24–26]. As a consequence, these populations represent ‘natural mapping experiments’ that can be studied to understand the relative contributions of the suite of alleles known to exhibit strong signatures of selection. These eight meiosis loci displaying highly differentiated alleles in *A. arenosa* were also reported in *A. lyrata* autotetraploids, along with signatures of extensive bidirectional gene flow [25]. At meiotic metaphase I in *A. arenosa*, chiasma frequency was reduced in autotetraploids carrying the derived alleles compared to the diploids, indicating an ongoing adaptive consequence of their evolution [8]. However, thus far, no formal confirmation of a direct effect on meiotic stabilisation in tetraploid *A. arenosa* or *A. lyrata* has been attributed to these evolved alleles.

In this study we fuse genomic, genetic and cytological approaches to investigate the effects of rapidly evolved adaptive haplotypes in these meiosis genes under strong selection. We measure the consequences of alternative evolved haplotypes at these loci in autotetraploid *A. lyrata, A. arenosa,* and natural introgressants of these species across a hybrid zone. Our analysis reveals functional evidence of a novel *ASY3* haplotype that modulates meiotic recombination in both *A. lyrata* and *A. arenosa* autotetraploids, thus stabilising chromosome bivalent formation and genomic integrity.

## Results

### A metaphase 1 analysis to determine meiotic stability in A. lyrata/A. arenosa populations

A thorough examination of meiotic stability was performed on fifty-two plants obtained from individual maternal seed lines sampled from tetraploid populations covering a range of known genomic backgrounds and demographic histories from the Wachau region and east Austrian Forealps. Relative genomic contributions (proportions of admixture) from *A. lyrata* and *A. arenosa* from sampled populations had previously been determined [25]. Sampled populations were: LIC, MOD, PIL, SCB, KAG, ROK (*A. lyrata* and *A. lyrata*-like hybrids, that contain >50% genomic contribution from *A. lyrata*) and, TBG, SEN and WEK (*A. arenosa* dominant/*A. lyrata* introgressants, that contain >50% genomic contribution from *A. arenosa*) (Fig 1). Meiotic stability was assessed in individual plants by performing cytological analyses on metaphase I (MI) chromosome spreads of pollen mother cells (PMCs). A rod bivalent forms when only one chiasma (the cytological manifestation of a CO) connects a homologous chromosome pair (Fig 2A, D1-3). A ring bivalent forms when chiasmata occur in both chromosome arms of homologous pairs (Fig 2A, D4). Quadrivalents are structures formed of four chromosomes, usually two pairs of homologous chromosomes and, multivalents form between multiple chromosomes either by chiasmata or interlocks (Fig 2C). As a control, chiasma frequency and distribution was scored at MI in diploid *A. lyrata* PMCs (Fig 2A). For the tetraploids, MI nuclei were scored as stable when 16 individual bivalents could be observed aligned on the MI plate and unstable if quadrivalents or multivalents were observed (Fig 2B-2C and S1 Fig). For each maternal line we scored blind the percentage of stable versus unstable nuclei, revealing a range from 20-100% (Fig 2E). Furthermore, we scored chiasmata as distal, interstitial or proximal to the centromere based on chromosome bivalent shapes (Fig 2D1-4). In unstable nuclei, only a proportion of individual bivalents could be scored per nucleus (ranging from 1-11 per cell), that were not associated with other chromosomes (Fig 2C, dashed ellipse). A FISH analysis utilizing the 5S and 45S rDNA probes revealed that in the samples of unstable nuclei all chromosomes that could be scored were observed both associated with multivalents and unassociated with multivalents. From the total sixteen pairs of chromosomes per nucleus, seven did not hybridize with the 5S and 45S rDNA probes, five with the 5S only, two with 45S only, and two with both 5S and 45S. However, the FISH analysis revealed that there was a chromosome bias for those that could be scored in the unstable nuclei with an underrepresentation of chromosomes containing the 45S rDNA nucleolar organizing regions (NOR) and an overrepresentation of chromosomes without the 45S rDNA. The expected frequency for bivalents that could be scored from the seventy seven chromosome counts based on a random expectation of occurrence from the proportion of labelled/unlabelled chromosomes was 34, 24, 9.5 and 9.5 for no probes, 5S only, 45S only, and both 5S and 45S, respectively, and the observed values were 46, 21, 6 and 4. A Chi-squared test revealed that these values were significantly different (χ[3]^2^ =9.54, *P* < 0.05), indicating that chromosomes with the 45S rDNA NOR were more likely to form multivalents (Fig 2C). As all chromosomes that could be scored were present in the sample from unstable nuclei, these were randomly grouped into pseudo-nuclei containing sixteen bivalents to determine chiasmata frequency and distribution, although with the caveat that these were less likely to form multivalents. Overall, significantly more chiasmata were observed in diploid *A. lyrata* MI bivalents than those from tetraploid stable or unstable nuclei (1.52±0.3, n=312; versus 1.12±0.2, n=960 and 1.26±0.3, n=590, respectively, Mann Whitney Test, P < 0.001). The frequency of distal chiasmata was not significantly different between bivalents in diploid and stable tetraploid nuclei, but was reduced in bivalents from unstable tetraploid nuclei (Fig 2F). Bivalents from stable tetraploid nuclei had significantly fewer interstitial and proximal chiasmata compared to bivalents from diploids and those from unstable nuclei, whereas interstitial and proximal chiasmata were not significantly different between bivalents from diploids and unstable tetraploid nuclei (Fig 2F).

**Fig 1.**
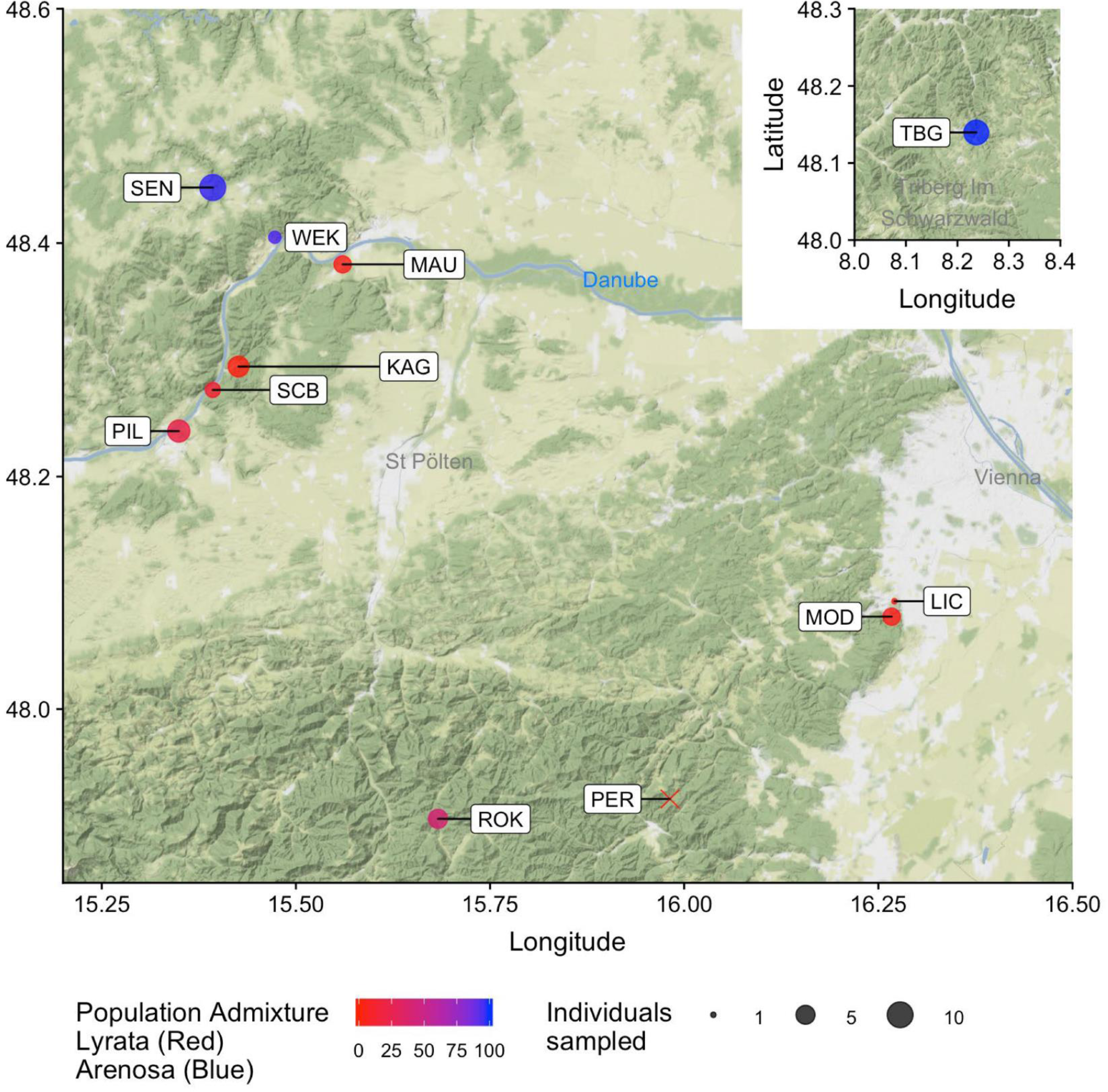
Map of maternal seed lines collection sites from the eastern Austrian fore Alps and Wachau valley. Collection sites are indicated as circles for tetraploid plants and an X for diploids. Circle size represents the numbers of plants sampled and analysed from each site and the shading from red to blue indicates the relative amounts of admixture between *A. lyrata* and *A. arenosa* populations determined in [25].

**Fig 2.**
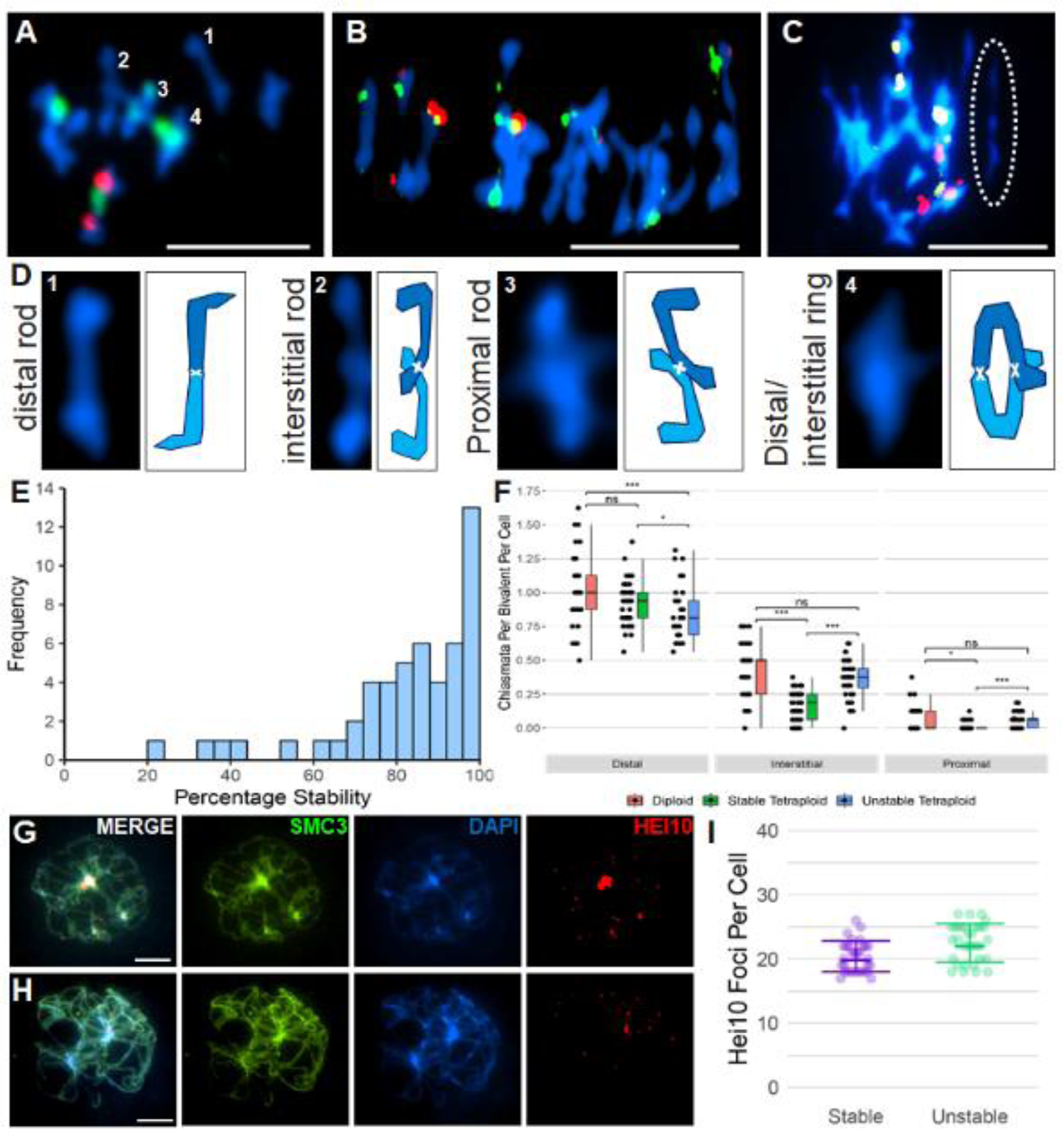
Cytological analysis of meiotic metaphase 1 stability, chiasma frequency and distribution. (A) Diploid *A. lyrata* bivalents stained with DAPI (blue) 5S rDNA (red) and 45S rDNA (green). (B) Stable autotetraploid meiotic metaphase I, stained with DAPI (blue) 5S rDNA (red) and 45S rDNA (green). (C) Unstable autotetraploid meiotic metaphase I, stained with DAPI (blue) 5S rDNA (red) and 45S rDNA (green) exhibiting one multivalent and one bivalent highlighted by the ellipse with white dashes that can be scored. (D) Four bivalents from A, with diagrammatic representations of homologous chromosomes in different shades of blue physically linked by chiasmata, shown by white crosses. (E) Histogram of the frequency of meiotic stability in individual plants from all populations. (F) Comparison of chiasmata position and frequency of *A. lyrata* bivalents from diploid (red), stable tetraploid (green) and, unstable tetraploid (blue). (G) Representative image of HEI10 foci at late pachytene in a stable nucleus, immunolocalised with SMC3 and counterstained with DAPI. (H) Representative image of HEI10 foci at late pachytene in an unstable nucleus, immunolocalised with SMC3 and counterstained with DAPI. (I) HEI10 counts from stable and unstable pachytene nuclei. Bars for A-C, G and H = 10µm.

A HEI10 immunocytological analysis was performed at late pachytene to confirm whether there was a difference in CO frequency between stable and unstable nuclei (Fig 2G-I). MAU8.11 (98% stability) and SEN2.2 (21% stability) were selected as extreme examples of meiotic stability (S1 Table). During pachytene HEI10 marks class I CO sites [27] and in our analysis HEI10 also marked heterochromatic DNA, which was not scored as designated class I CO sites. In stable MAU8.11 nuclei, an average of 20.4 HEI10 foci per pachytene (n=30) were scored and in the unstable SEN2.2 nuclei, an average of 22.5 HEI10 (n=30) were scored, revealing that the unstable nuclei had significantly greater numbers of HEI10 foci (Wilcoxon rank sum test, P<0.05) (Fig 2I). Stable nuclei contained an average of 1.28 HEI10 foci/bivalent, whereas the unstable nuclei contained an average of 1.4 HEI10 foci/bivalent.

### *Association of haplotypes with meiotic stability in* A. lyrata/A. arenosa *tetraploids*

The fifty-two tetraploid plants phenotyped for male meiotic stability were then genotyped for the proportion of each meiosis gene haplotype by high-throughput sequencing. Accurate genotyping required obtaining precise population reference sequences from published genomic data from these populations [25]. Degenerate primers were used to amplify full length gene amplicons of the eight meiosis genes from the fifty-two tetraploid plants (including exons and introns) for construction of Nextera LITE libraries. Libraries were barcoded per plant and sequenced by MiSeq, generating an average sequence depth across all loci of >2000x, from which we determined the proportion of each haplotype per plant by SNP frequency. Because the coding regions of all eight meiosis genes from representative populations were cloned and Sanger sequenced (described below in ‘Adaptive polymorphisms in meiosis genes’), it was possible to resolve individual haplotypes. As these plants were drawn from a diversity of wild populations it was not surprising that an average of 7.25 alleles were identified for each of the eight meiosis genes. Given the limited sample size of fifty-two individuals it was not possible to statistically interrogate associations between all individual haplotypes for the eight genes. Consequently, similar haplotypes were collapsed together and classified into two groups: haplotypes with derived tetraploid alleles, and those with ancestral diploid alleles. Derived tetraploid haplotypes were those possessing conserved polymorphisms compared to the diploid reference sequences. For each gene (except ZYP1b, which was homozygous in all populations tested) a large proportion of the individuals carried four derived tetraploid haplotypes, whilst the others carried a mixture of derived tetraploid and diploid haplotypes. We therefore tested whether the presence of diploid haplotypes influenced meiotic stability. To do this, we classified the allele state at each of the eight meiosis genes in each individual tetraploid as either homozygous (i.e. exclusively either derived tetraploid haplotypes, or alternatively ancestral diploid haplotypes) or heterozygous (individuals harbouring both ancestral and derived alleles together at a given locus), and tested for any associations between these genotypes and meiotic stabilty by cytological analysis (Bonferroni corrected pairwise Mann-Whitney-Wilcoxon; S1 Table). This revealed that only the meiotic chromosome axis gene *Asynapsis3* (*ASY3)* had a significant effect on meiotic stability (Fig 3). Plants that were heterozygous for the *ASY3* ancestral diploid haplotype and the derived tetraploid haplotype had significantly more unstable male metaphase I nuclei than plants homozygous for the derived ASY3 haplotype (4n Hom 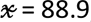, IQR = 15.1, n= 41, 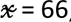, IQR =41.7, n=11, p=0.008). There was a large range of meiotic stability within the *ASY3* heterozygotes, so smaller effects from the other seven meiosis genes cannot be excluded. For example SMC3 showed a trend, whereby the ancestral diploid allele may be associated with lower meiotic stability than the tetraploid homozygotes or diploid/tetraploid allele heterozygotes, although sampling sizes were not great enough to statistically confirm this trend.

**Fig 3.**
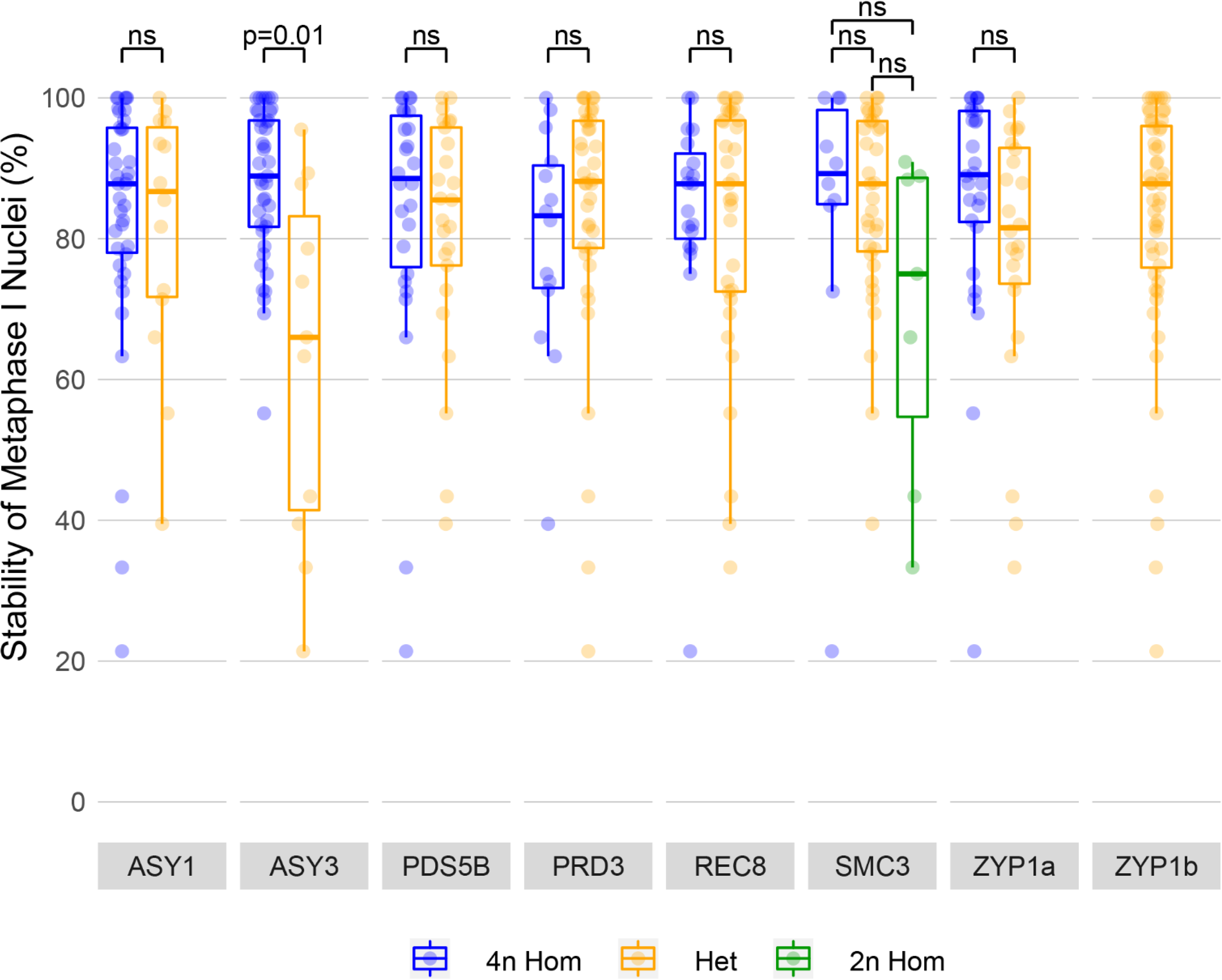
Association analysis of meiotic chromosome stability at metaphase I with meiosis gene haplotypes. Individual plants are collapsed into groups containing either homozygous 4n derived (blue), 2n/4n heterozygous (orange) or 2n homozygous ancestral (green) haplotypes from all populations tested. Each data point represents the male meiotic stability of an individual plant for each specific haplotype.

### Adaptive polymorphisms in meiosis genes

Previous studies have inferred polymorphic amino acids in meiosis genes between diploids and tetraploids in *A. arenosa* and *A. lyrata* by aligning short read sequences to the *A. lyrata* reference, but could not infer contiguous autotetraploid alleles [8, 25, 28]. To overcome this and resolve individual haplotypes, we amplified, cloned and Sanger sequenced the coding regions of the eight meiosis genes from diploid *A. arenosa* SNO and *A. lyrata* PER populations, tetraploid *A. arenosa*, SEN, TBG and WEK populations, as well as *A. lyrata* KAG and MAU populations. This approach provided high-resolution sequence polymorphism data for a total of three hundred and twenty cDNA transcripts, consisting of fifty-eight alleles with an average of seven alleles for each of the eight meiosis genes, thus allowing us to detect both structural variation including indels and divergent SNP variation (Fig S2, S2-4 Tables). The eight encoded proteins associated with the synaptonemal complex are conserved at the secondary and tertiary amino acid level, rather than at the primary sequence [29]. It is therefore difficult to infer functional or non-functional amino acid polymorphisms with complete confidence, but results from KinasePhos2.0 and NetPhos3.1 suggest that overall 45% of those we detect were either loss or gain of putative serine/threonine phosphosites and 55% were non-phosphosites. The ASY3 derived adaptive allele and ZYP1b exhibited the greatest quantity of residue changes compared to the diploids that were conserved in tetraploid populations (45 and 44, respectively) (SFig 4), whilst SMC3 had none (although there were population specific SMC3 SNPs). Of the 45 polymorphic residues between the ancestral *A. lyrata* diploid ASY3 allele and the derived tetraploid ASY3 allele, 27 were due to a tandem duplication (TD allele) in a serine-rich region of the protein. The serine-rich region is upstream of the coiled-coil domain, possessing putative ATM/CKII phosphosites and a predicted SUMO site (K556, GPS-SUMO), and the TD allele contains 14 serines in this region, compared to 7 in the ancestral *A. lyrata* non-duplicated allele (ND) (Fig 4A). As the *ASY3 TD* allele had highest sequence similarity to the diploid *A. lyrata ASY3 ND* allele, to investigate the provenance of the adaptive tandem duplication, we exhaustively screened the local diploid *A. lyrata* population (PER) geographically adjacent to the *A. lyrata* autotetraploid LIC and MOD populations (Fig 1). One hundred and twenty-eight plants from the diploid *A. lyrata* PER population were screened, but the *ASY3 TD* allele was not identified, indicating the current absence (or vanishingly low frequency) of the *TD* allele, although it did reveal the presence of a deletion (DEL) allele at 7% frequency, where the entire serine-rich region is absent. We therefore cloned and sequenced genomic DNA from *ASY3 ND*, *TD* and *DEL* and aligned them. This revealed a 78bp region of exon 2 is duplicated in-frame in the *ASY3 TD* allele that is missing in the *ASY3 DEL* allele between two AGAGA sites that possess DNA microhomology. We speculate that this DNA microhomology may have been instrumental in the formation of the *ASY3 TD* allele by homologous DNA repair through a replication error or during meiotic recombination (Fig 4B).

**Fig. 4.**
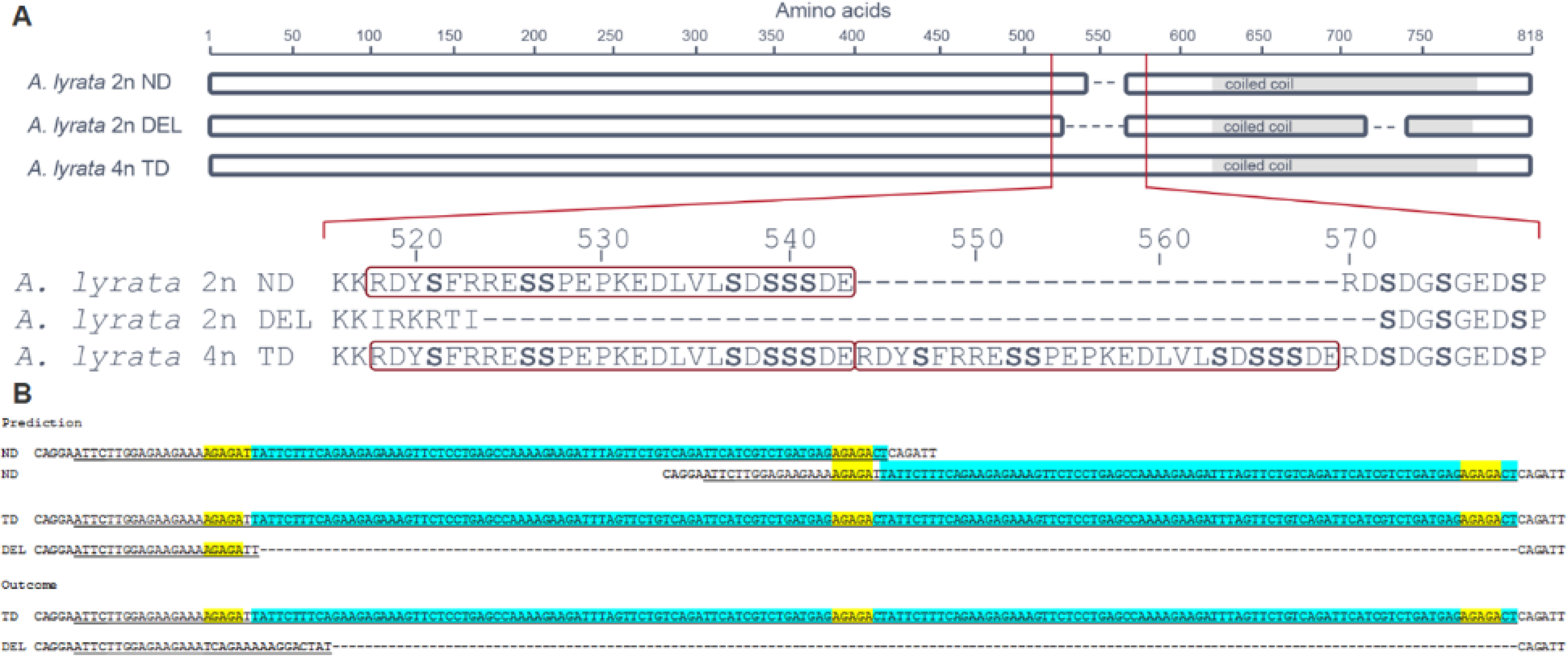
Structural variants of the three major *A. lyrata ASY3* alleles. (A) The serine-rich region (red box) with serines highlighted in bold in the non-duplicated (ND) ancestral diploid allele, absence in the deletion (*DEL*) diploid allele and the tandemly duplicated (TD) derived tetraploid allele possessing two serine-rich regions (red boxes) and putative SUMO sites at K517, K531 and K556. (B) Major structural variation at the ASY3 locus suggesting that DNA microhomology (highlighted yellow) between the ancestral alleles may have led to the formation of the *TD* allele (underline=exon, blue=region of duplication).

### Meiosis gene flow between A. lyrata/A. arenosa tetraploid populations

We categorized the admixture proportions of the tetraploid populations into *A. lyrata* dominant (LIC, MOD, SCB, KAG, PIL and ROK) or *A. arenosa* dominant (WEK, SEN and TBG), based on a demographic analysis [25]. The Sanger sequenced coding regions of the eight meiosis genes provided high quality reference data to determine allelic origin and to infer the direction of gene flow. Maximum likelihood and Bayesian phylogenetic analyses showed that for each gene these sequences clustered cleanly according to ploidy (S5-6 Fig). The ancestral diploid sequences were further divided into *A. lyrata* or *A. arenosa.* Tetraploid sequences clustered into separate alleles irrespective of species, although population differences persisted (S5-6 Fig). The *ASY3 TD* allele that associates with meiotic stability clusters with the *A. lyrata* diploid sequence (S5B Fig), as does *PDS5b* (S5C Fig). Conversely, *ASY1, PRD3, REC8, SMC3* and *ZYP1a/b* had highest homology with diploid *A. arenosa* (S5A, B, D-H Fig). The adaptive *A. lyrata ASY3 TD* allele is present at 99% frequency in tetraploid *A. arenosa* populations tested and 95% in tetraploid *A. lyrata* populations (Fig 5). The analysis revealed a small number of ancestral diploid *A*. *arenosa* and *A*. *lyrata* alleles in these populations, except *ZYP1b* that was completely homozygous. In contrast, the *ASY1* allele derived from diploid *A. arenosa* had a frequency of 94% in *A. arenosa* tetraploid and 93% in *A. lyrata* tetraploid, indicating bidirectional gene flow (Fig 5), as well as bidirectional gene flow of ancestral *SMC3* and *REC8* diploid alleles in the tetraploids (Fig 5).

**Fig 5.**
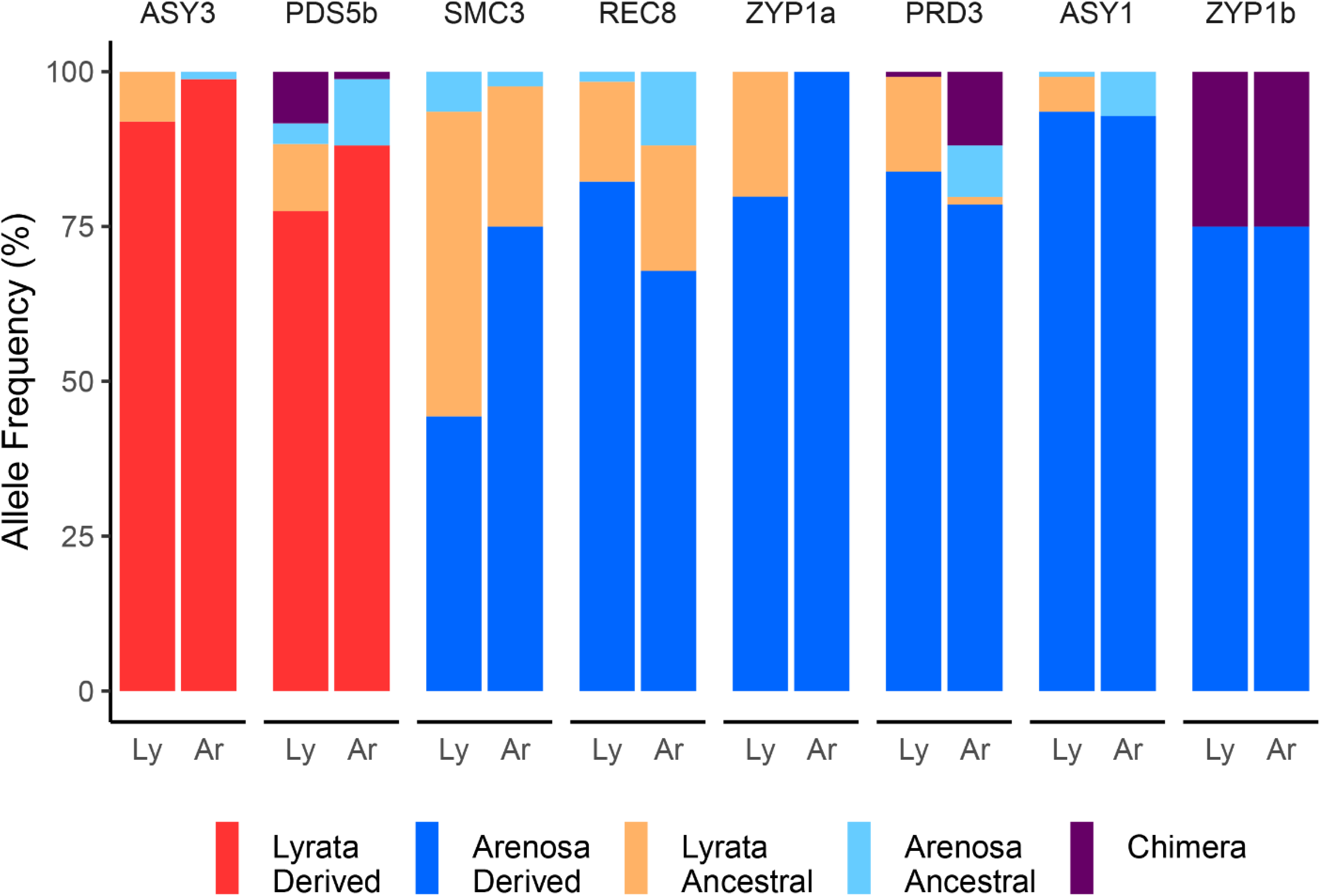
Contrasting origins and polymorphisms of adaptive meiosis gene haplotypes in tetraploid introgressants of *A. lyrata*/*A. arenosa*. Haplotype frequencies of adaptive meiosis alleles from *A. lyrata* (*Ly*) and *A. arenosa* (*Ar*) populations.

### Widespread chimerism in meiosis gene alleles

Our analysis of Sanger and MiSeq data identified a novel chimeric allele of *ZYP1b* in all tetraploid populations; chimeric *PRD3* in SCB, SEN and WEK; and chimeric *PDS5b* in ROK. While rare chimeras could potentially result from PCR artefacts [30], these alleles have the same breakpoints in multiple individuals which suggests alterations in the genomic DNA through homologous DNA repair *in planta* [31]. At the 3’ end of all *ZYP1b* tetraploid alleles we detect evidence for a 474bp gene conversion (GC) to *ZYP1a* (S7A Fig). There is also evidence of GC in *PRD3* between *A. arenosa* and *A. lyrata* ancestral diploid alleles (S7B Fig). In eleven plants from *A. arenosa* and *A. lyrata* populations, the first 740bp of the diploid *PRD3* allele is more similar to the diploid *A. arenosa* than to *A. lyrata* (7 vs 21 SNPs), while the remaining 625bp of coding sequence has a higher similarity to diploid *A. lyrata* than to *A. arenosa* (5 vs 24 SNPs). In *PDS5b,* at the 5’ end of tetraploid alleles from five *A. arenosa* plants we observe evidence for a GC of the first two exons to the diploid allele, providing evidence of GC (or CO) between ploidy levels (S8A Fig). In addition, analysis of genome resequencing data [25] revealed a 3’ GC from a diploid *ASY1 A. lyrata* allele to a tetraploid *A. arenosa* allele in the KAG population (S8B Fig). The widespread presence of such evidence of gene conversion products in these loci exhibiting the most dramatic signatures of selection suggests a mechanism by which the peaks of differentiation generally found in this system are so narrow [3, 8, 25, 32, 33], despite a recent origin, and possible bottleneck, of the tetraploids [25].

## Discussion

Here we aimed to determine the impact of strongly selected meiosis alleles that underwent recent selective sweeps on the rapid evolution of autotetraploid meiotic stability in *A. lyrata/A. arenosa hybrids* and introgressants and to trace their evolutionary origin. By associating genotypic and cytological phenotypic data we provide evidence that *ASY3* is the major locus currently stabilising autotetraploid male meiosis in these populations. We identified structural variation of meiosis alleles including a novel derived, *ASY3* allele with a tandem duplication (TD) in a serine-rich region that underlies the stable chromosome meiotic phenotype in the tetraploids, as well as novel *ASY1, PDS5b, PRD3* and *ZYP1b* chimeric alleles between diploids and tetraploids and *A. arenosa* and *A. lyrata* origins.

A cytological metaphase I (MI) analysis revealed that chiasmata in the autotetraploids were significantly reduced in both stable and unstable nuclei compared to diploid *A. lyrata*. Moreover, chiasma frequencies in meiotically stable nuclei were significantly reduced in regions proximal and interstitial to the centromere. A shift in chiasma distribution may reflect a fundamental mechanism for meiotic adaptation to autopolyploidy [34]. The unstable nuclei occur due to unregulated meiotic recombination between multiple chromosomes, either homologous or non-homologous. All chromosomes appeared to associate with multivalents, although there was a bias for the 45S rDNA containing chromosomes, which may be due to the NORs clustering during prophase I, thus bringing non-homologous chromosomes into close proximity. A reduction in numbers of chiasmata in stable nuclei was supported by an immunocytological approach counting HEI10 foci that mark designated class I COs. HEI10 foci numbers were significantly lower in the stable pachytene nuclei (20.4 HEI10 foci per cell) compared to the unstable nuclei (22.5 HEI10 foci per cell)(P<0.01).

Our genotype-phenotype association study revealed that among these eight meiosis genes, the allele state of the structurally variable meiotic chromosome axis protein ASY3 was the major factor governing whether nuclei were stable. We hypothesise that the ancestral diploid *ASY3 ND* allele promotes high levels of proximal and interstitial chiasmata, but in the tetraploid it acts dominantly over the evolved *ASY3 TD* allele, promoting interstitial and proximal chiasmata as well as complex chromosome structures including multivalents. Such multivalents have previously been observed in diploid *A. thaliana ZYP1^RNAi^* lines where the authors postulated that chiasmata may have formed between extensive duplications with high sequence similarity on non-homologous chromosomes [23]. The tandemly duplicated serine-rich region in the ASY3 TD allele may function in a manner similar to the budding yeast N-terminus MSH4 degron that destabilizes the protein until it is phosphorylated [35]. Further studies are required to determine if the serine-rich region destabilizes the ASY3 protein thus creating a hypomorphic variant or whether these sites are phosphorylated. The analysed brassica SC phosphoproteome [36] did not recover peptides for ASY3 in the serine-rich region, although similar (serine-aspartic acid) residues recovered from ASY1 were phosphorylated. The ASY3 TD allele also contains 19 derived residues outside the serine-rich region, of which 10 are predicted phosphosite gains or losses, that cannot be ruled out as functionally important along with unknown *trans* effects. Chiasmata are distalized in *A. thaliana* chromosome axis mutants *asy1, asy3* and *asy4*, presumably due to telomere proximity enabling sufficient inter-homolog pairing, whereas a complete meiotic axis is required to promote high levels of recombination between spatially separated regions in nuclei along the arms of the chromosomes [19, 20, 37].

From our data and extensive literature we can speculate a model wherein the serine-rich duplicated region destabilizes the ASY3 TD protein and may target the protein for degradation by the proteasome [35, 38–40]. A destabilized protein may be hypomorphic to the ancestral ASY3 ND allele as there less protein may be available to bind to the chromosome axes and/or it could be less effective at promoting interhomolog recombination when bound at the axis. A hypomorphic protein may act similarly to ASY1, ASY3 and ASY4 mutants in distalizing chiasmata [19, 20, 37]. However, when heterozygous with the dominant ancestral ASY3 ND allele, rates of inter-homolog and non-homologous recombination increase. We hypothesise that axis components that favour interstitial and proximal recombination in diploids promote associations with non-homologous chromosomes in the tetraploids, especially in regions with high sequence homology. However, for stable bivalents, once a distal CO site is designated, CO interference may prevent further COs forming. In budding yeast, the TopoII interference pathway requires SUMOylation of TopoII and Red1, the ASY3 orthologue [22, 41]. The serine-rich duplicated region in ASY3 possesses a predicted SUMO site, which could play a role in protein function, although this hypothesis needs functional testing.

Phylogenetic analysis revealed that the *ASY3 TD* allele most likely originated from diploid *A. lyrata*, consistent with [25]. A screen of the diploid *A. lyrata* population geographically adjacent to the tetraploids for the *ASY3 TD* did not detect the allele, but instead identified one with the same region deleted (*DEL*). The *ASY3 DEL* allele coding region is in-frame and contains a second deletion in the coiled-coil. It may be a coincidence that the same region is lost and gained in the *ASY3* alleles, or that this sequence is more susceptible to DNA replicative errors. The region of interest is flanked by DNA microhomology (AGAGA) that is positioned at the putative exchange points and may have led to DNA polymerase slippage or aberrant replication fork repair. It is unlikely to have occurred during meiotic recombination due to the low level of sequence homology. However, we observe numerous examples of apparent gene conversion that appear to have arisen during meiotic recombination in *ASY1, PRD3, PDS5b* and *ZYP1a/b* between both species and ploidies. The bidirectional gene flow between *A. arenosa* and *A. lyrata* tetraploids has enlarged the gene pool for beneficial alleles to be borrowed and selected upon and the novel gene converted chimeric alleles may precisely coalesce advantageous sequences from differing origins for adaptation, although this would require further testing with an even larger sample size.

Evidence suggests that the origin of the adaptive *ASY3 TD* allele in the tetraploid populations is relatively recent, but as it is under extreme selection, it has spread extensively and introgressed within tight boundaries in the *A. lyrata*/*A. arenosa* hybrid genomes tested [25]. Our current analysis has provided haplotype-level sequence data that supports this hypothesis. The orthologous diploid ASY3 alleles from *A. arenosa* and *A. lyrata* are highly divergent, and yet the tetraploid *ASY3 TD* allele is very similar among tetraploid populations. We speculate that is therefore possible that preceding the origin of the *ASY3 TD* allele, gene flow of adaptive alleles from *A. arenosa* (*ASY1, PRD3, REC8, SMC3* and *ZYP1a/b*) was necessary to establish meiotic stability in the nascent *A. lyrata* tetraploids, but has since been relaxed due to the presence of the predominant *ASY3 TD* allele.

The *A. lyrata* autotetraploid populations contain plants with variable levels of meiotic stability as well as relatively high frequencies of diploid-like alleles. The diploid alleles may persist due to: a) continuous gene flow via unreduced gametes [42]; b) being beneficial in certain environmental conditions e.g. high altitudes [43]; c) advantageous during female meiosis (COs are reduced in *A. thaliana* female compared to male [44]; d) in the case of SMC3, diploid alleles may be beneficial for male meiosis; e) inability to purge genetic load in autotetraploids [45] and; f) limited effect on overall male pollen fecundity, due to an excess of grains transmitted, despite variable quality.

Taken together, our data indicate multiple mechanisms for rapid meiotic evolution in autotetraploid *A. lyrata*. They reveal a predominant association of a duplication of the serine-rich region in the ASY3 TD allele with MI stability. Furthermore, tetraploid *A. lyrata* has introgressed *ASY1 PRD3, REC8, SMC3* and *ZYP1a/b* alleles from *A. arenosa* by gene flow. Finally, novel chimeric genes of *ASY1, PDS5b, PRD3,* and *ZYP1a/b* have arisen evidently through gene conversion, suggesting highly dynamic mechanisms to generate variation that may be selected upon by evolution to ensure meiotic success in these populations.

## Material and Methods

### Cloning and sequencing of meiosis gene transcripts

Plants were grown from seed to obtain fresh flower buds from diploid *A. arenosa* SNO and *A. lyrata* PER populations, and tetraploid *A. arenosa*, SEN, TBG and WEK and *A. lyrata* KAG and MAU populations [25]. These buds were collected, flash frozen in liquid nitrogen and stored at -80C until RNA extraction. Total RNA was extracted using a Bioline ISOLATE II RNA Plant Kit (Bioline Ltd, London, UK), following manufacturer’s instructions, eluting into a final volume of 60 μl nuclease free water. Concentration and purity were determined using a NanoDrop spectrophotometer (LabTech International, Lewes, UK) and one microgram of total RNA was electrophoresed on a non-denaturing 1% (w/v) agarose gel to check for degradation. First strand cDNA was reverse transcribed from 0.5 μg of total RNA using a Quantitect Reverse Transcription Kit (Qiagen, Hilden, Germany) that incorporates genomic DNA removal prior to reverse transcription. The coding regions of 8 meiosis genes (*ASY1*, *ASY3*, *PRD3*, *PDS5b*, *REC8*, *SMC3*, *ZYP1a* and *ZYP1b*) were amplified by PCR using 0.2 μM primers (STable 5) designed using HiSeq data [33] and Platinum^™^ *Taq* DNA Polymerase High Fidelity (ThermoFisher Scientific, MA, USA). PCR conditions were as follows: 94 °C for 2 min, followed by 35 cycles of 94 °C for 15 s, 60-65 °C for 30 s and 68 °C for 2-5 min (see S5 Table), with a final extension of 68 °C for 5-10 min. PCR products were electrophoresed on a 2% (w/v) agarose gel, and single bands of the expected size were excised and purified with a Monarch^®^ DNA Gel Extraction kit (New England Biolabs, MA, USA). Purified PCR products were cloned into pCR-XL-TOPO^™^ vector using a TOPO^™^ XL PCR Cloning Kit following the manufacturer’s instructions. For each gene a total of 8 clones from each plant were isolated from overnight LB cultures using an ISOLATE II Plasmid Mini Kit (Bioline) prior to sequencing with universal M13F and M13R primers by GATC Biotech (Konstanz, Germany). Nucleotide sequences of the cDNAs were processed in Geneious 11.1.2 (https://www.geneious.com) to remove vector and low-quality sequence before using BLASTN to search the June 2010 (v.1.0/INSDC) assembly of the North American *A. lyrata* reference genome [46] and NCBI nonredundant (nr) database for confirmation that the obtained cDNAs were the expected gene transcripts. Primer walking was then used to sequence the entire length of the transcript. For each meiosis gene, cDNAs from each population were aligned with the respective Ensembl gene predictions from the *A. lyrata* reference genome (S6 Table), and to act as outgroups, *A. thaliana* transcripts obtained from The Arabidopsis Information Resource (TAIR) using the MUSCLE 3.8.425 plugin in Geneious 11.1.2 with default settings [47]. Phylogenetic trees were constructed using a maximum likelihood (ML) method with PhyML v3.3 (Guindon et al, 2010) and bootstrap testing (1000 replicates). The best nucleotide substitution method was determined with the Find Best-Fit Substitution Model in Mega v10.0 (Kumar et al, 2018). The ML phylogenetic trees were confirmed by a Bayesian approach with MrBayes v3.2.6 (Huelsenbeck and Ronquist, 2001) and bootstrap testing of 1000 replicates.

### Meiotic haplotype genotyping

Genomic DNA was extracted from leaf material of each of the 52 plants in the study using a DNeasy Plant Mini Kit (Qiagen) and eluting into 100 μl nuclease free water. Full length coding regions of each of the 8 meiosis genes (including introns) were amplified from this genomic DNA by PCR using Platinum^™^ SuperFi^™^ Green PCR Master Mix (ThermoFisher Scientific) and 0.5 μM primers (S5 Table) designed against the Sanger sequenced cDNA of the 8 meiosis genes (see above). PCR conditions were as follows: 98 °C for 30 s, followed by 35 cycles of 98 °C for 30 s, 60-63 °C for 10 s and 72 °C for 2.5-10 min (see S5 Table), with a final extension of 72 °C for 5-10 min. PCR products were electrophoresed on a 1% (w/v) agarose gel, and single bands of the expected size were excised and purified with a Monarch DNA Gel Extraction kit (New England Biolabs). Libraries were constructed using 1ng of input DNA in a Low Input, Transposase Enabled (LITE) pipeline developed at the Earlham Institute (Norwich, UK) and based on the Illumina Nextera kits (Illumina, San Diego, CA, USA) [48]. Each library was constructed using unique 9 bp dual index combinations allowing samples to be multiplexed. Pooled libraries were size selected between 600 and 750bp on a BluePippin (Sage Science, Beverly, MA, USA) 1.5% Cassette and then sequenced with a 2×250bp read metric on an Illumina MiSeq sequencer.

MiSeq fastq files were imported into Geneious 11.1.2 (https://www.geneious.com) and R1 and R2 reads paired. Quality trimming of reads was performed with the BBDuk Adaptor/Quality Trimming v.37.64 plugin with default settings (Min quality:20; Min overlap:20; Min length:20). For each of the 52 individuals, trimmed reads were mapped to each of the 8 meiosis genes (S6 Table) from the North American *A. lyrata* reference genome [46] using Geneious 11.1.2 Read Mapper (Medium sensitivity; 5 iterations and default settings). SNPs relative to the reference genome genes were then called (Minimum Variant Frequency 0.25; Maximum Variant P-value 6×10^-6^) and used to identify and determine the proportion of 2n and 4n alleles for each gene per plant using a set of allele specific SNPs (as revealed from the Sanger sequencing of meiosis gene transcripts described above; S7 Table). Allele specific indels (eg ASY3 TD) were identified by associated SNPs.

### Cloning and sequencing of ASY3 alleles

Genomic DNA was extracted from diploid *A. lyrata* PER plants as above and a short section of *ASY3* was PCR amplified using 0.5 μM primers (S5 Table) and MyTaq^™^ Red Mix (Bioline). PCR conditions were as follows: 95 °C for 2 min, followed by 35 cycles of 95 °C for 30 s, 69 °C for 30 s and 72 °C for 30 s, with a final extension of 72 °C for 5 mins. PCR products were electrophoresed on a 2% (w/v) agarose gel and the lower band (∼125 bp) corresponding to a partial *ASY3 DEL* allele was excised and purified with a Monarch^®^ DNA Gel Extraction kit (New England Biolabs). Purified PCR products were cloned into pCR^™^4-TOPO^®^ vector using a TOPO^™^ TA Cloning^™^ for Sequencing Kit (ThermoFisher Scientific) following manufacturer’s instructions. A total of 12 clones were isolated, purified and Sanger sequenced as above. Nucleotide sequences were processed in Geneious 11.1.2 (https://www.geneious.com) to remove vector and low-quality sequence before aligning with *ASY3 ND*/*TD* transcript alleles as described above. Primers designed against the partial *DEL* sequence were used to obtain the 3’ end of the transcript from 3 µg *ASY3 ND/DEL* heterozygous 2n *A. lyrata* (PER) floral bud total RNA using a GeneRacer^™^ Kit (ThermoFisher Scientific) following manufacturer’s instructions. Purified PCR products were cloned into pCR^™^4-TOPO^®^ vector, sequenced and processed as above.

*ASY3 ND*, *TD* and *DEL* alleles were PCR amplified from genomic DNA extracted from diploid and tetraploid *A. lyrata* respectively using 0.2 µM primers designed against *ASY3* cDNA sequences obtained above (S5 Table) and Q5^®^ High-Fidelity DNA Polymerase (New England Biolabs). PCR conditions were as follows: 98 °C for 2 min, followed by 35 cycles of 98 °C for 10 s, 63°C for 30 s and 72 °C for 4 min, with a final extension of 72 °C for 10 min. PCR products were electrophoresed on a 1% (w/v) agarose gel, and single bands of the expected size were excised and purified as above. Purified PCR products were cloned into pDrive (Qiagen) and sequenced by Eurofins Genomics (Ebersberg, Germany).

### Cytology

Chromosome spreads were performed [49] [25] on all populations used in this study. The HEI10 immunocytological analysis was performed using the protocol [50] with anti-AtSMC3 rat and anti-AtHEI10 rabbit antibodies described in [51]. Nikon Eclipse Ci and Ni-E microscopes installed with NIS Elements software were used to capture images of chromosomes.

### Protein predictions

Protein post-translational predictions were provided by KinasePhos2.0 [52] and NetPhos3.1 [53] and SUMO sites were predicted by GPS-SUMO [54].

### Statistical analyses and Map drawing

Statistical analysis was performed using the R *Stats* package. Mann-Whitney Wilcoxon tests were performed with function *wilcox.test*. Bonferoni adjusted p values were calculated using the function *p.adjust*. The map was drawn using ggmap [55].

### Data availability

All sequences in this study including cDNA transcripts and genomic DNA sequences have been deposited in the DDBJ/EMBL/GenBank databases under accession numbers MN512718 - MN513026 and MN520243 - MN520257. MiSeq amplicon reads have been deposited in the NCBI Sequence Read Archive (SRA; https://www.ncbi.nlm.nih.gov/sra) database under BioProject ID PRJNA575228.

## Acknowledgments

We would like to thank Neelam Dave and Maria Spezzati for providing technical support and Chris Franklin and Holger Puchta for helpful discussions during the project. DEEPSeq at the University of Nottingham provided MiSeq data.

## Funding

This work was funded by UKRI through Biological and Biotechnological Sciences Research Council (BBSRC) New Investigator grant (BB/M01973X/1) to JDH. This work was supported by the European Research Council (ERC) under the European Union’s Horizon 2020 research and innovation programme (ERC-StG 679056 HOTSPOT), via a grant to LY; and the BBSRC (BB/P013511/1), via a grant to the John Innes Centre (LY).

### Author contributions

PJS, CLG, DH, JDH performed lab work. PJS, MGF and JDH analysed data. JDH and LY conceived the project. PJS, MGF, DH, RS, LY and JDH wrote the manuscript.

## Supplemental Tables

**S1 Table.**
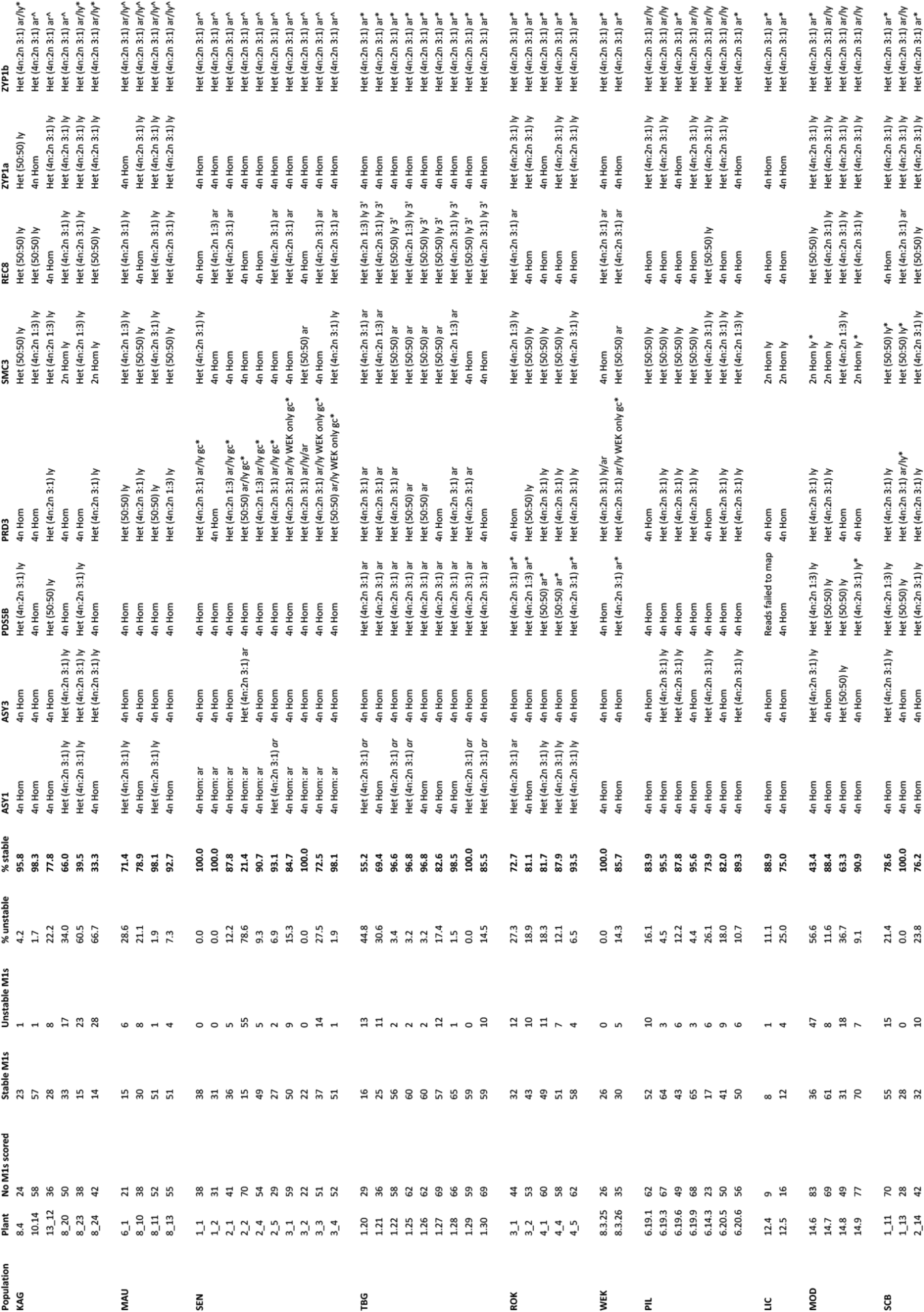
Genotype and phenotype data.

**S2 Table.**
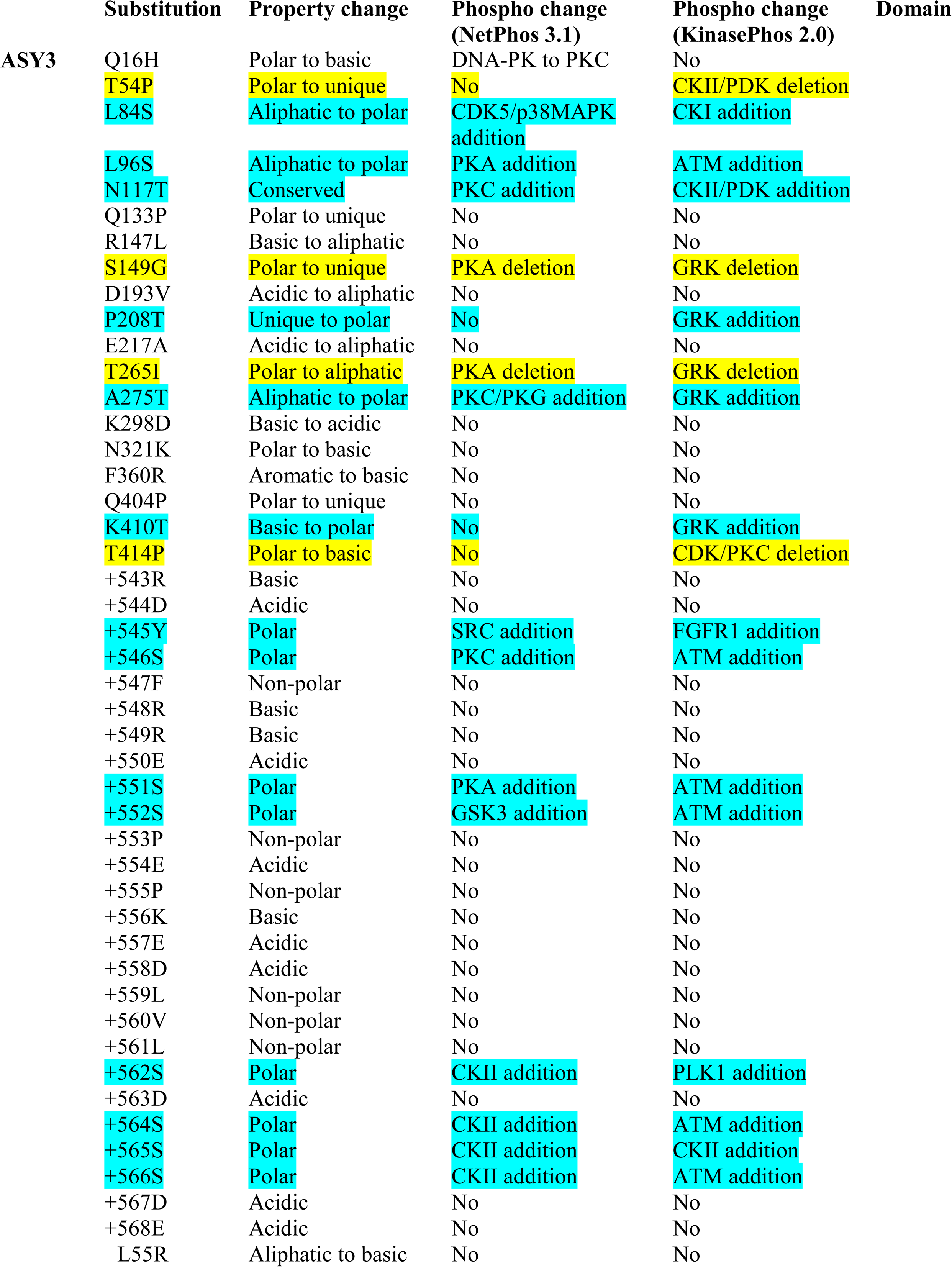
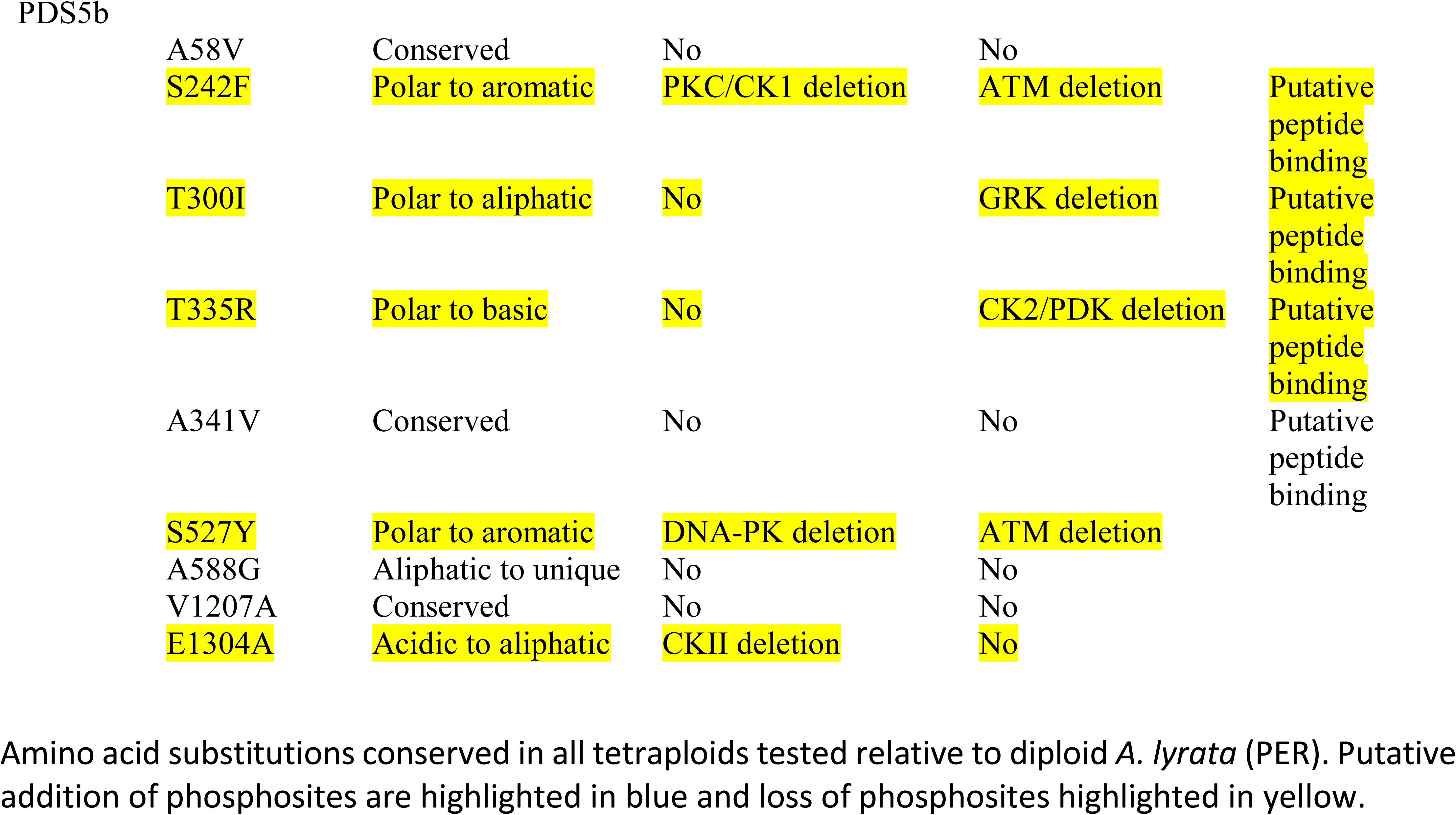

**S3 Table.**

| Substitution | Property change | Phospho change<br>(NetPhos 3.1) | Phospho change<br>(KinasePhos2.0) | Domain |
| --- | --- | --- | --- | --- |
| L84S | Aliphatic to polar | CDK5/p38MAPK addition | CK1 addition |  |
| R147L | Basic to aliphatic | No | No |  |
| S149G | Polar to unique | PKA deletion | GRK deletion |  |
| R162T | Basic to polar | No | PLK1 addition |  |
| L172Q | Aliphatic to polar | No | No |  |
| K298N | Basic to polar | CKII addition | No |  |
| N321K | Polar to basic | No | No |  |
| K342N | Basic to polar | No | No |  |
| A352T | Aliphatic to polar | PKC addition | PKA addition |  |
| F360R | Aromatic to basic | No | No |  |
| Q380H | Polar to basic | ATM to PKC | No |  |
| Q404P | Polar to unique | No | No |  |
| K410T | Basic to polar | No | GRK addition |  |
| T414P | Polar to unique | No | PKC/CDK deletion |  |
| A467V | Conserved | No | No |  |
| T478I | Polar to aliphatic | CKII deletion | PKC/CDK deletion |  |
| M482T | Aliphatic to polar | No | GRK addition |  |
| N494T | Conserved | DNA-PK deletion | GRK addition |  |
| S525R | Polar to basic | PKA to p38MAPK | ATM deletion |  |
| K530N | Basic to polar | No | No |  |
| S538P | Polar to unique | CKII deletion | ATM deletion |  |

Amino acid substitutions in ASY3 of 2n *A. arena* (SNO) relative to 2n *A. lyrata* (PER). Putative addition of phosphosites are highlighted in blue and loss of phosphosites highlighted in yellow.

**S4 Table.**
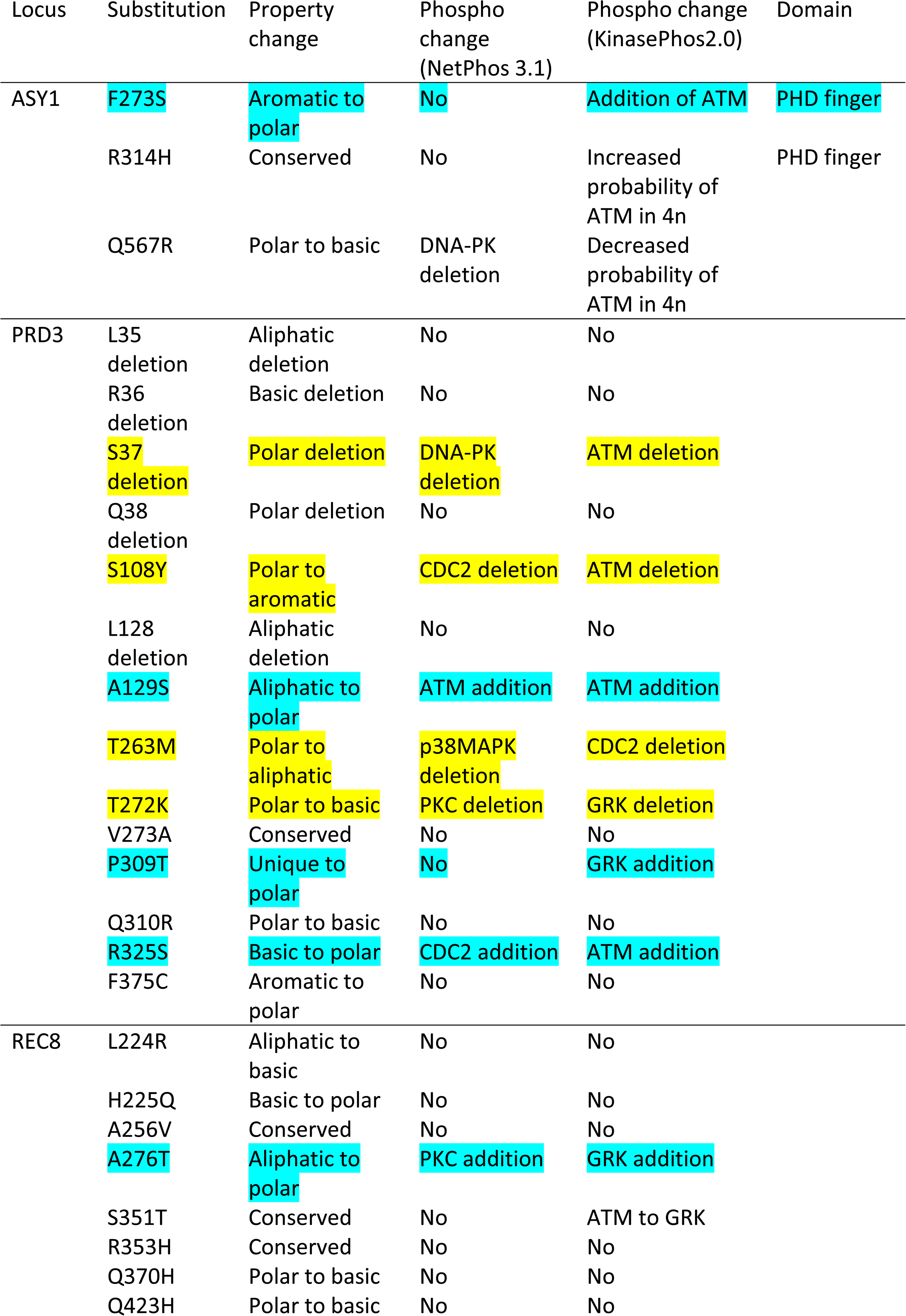
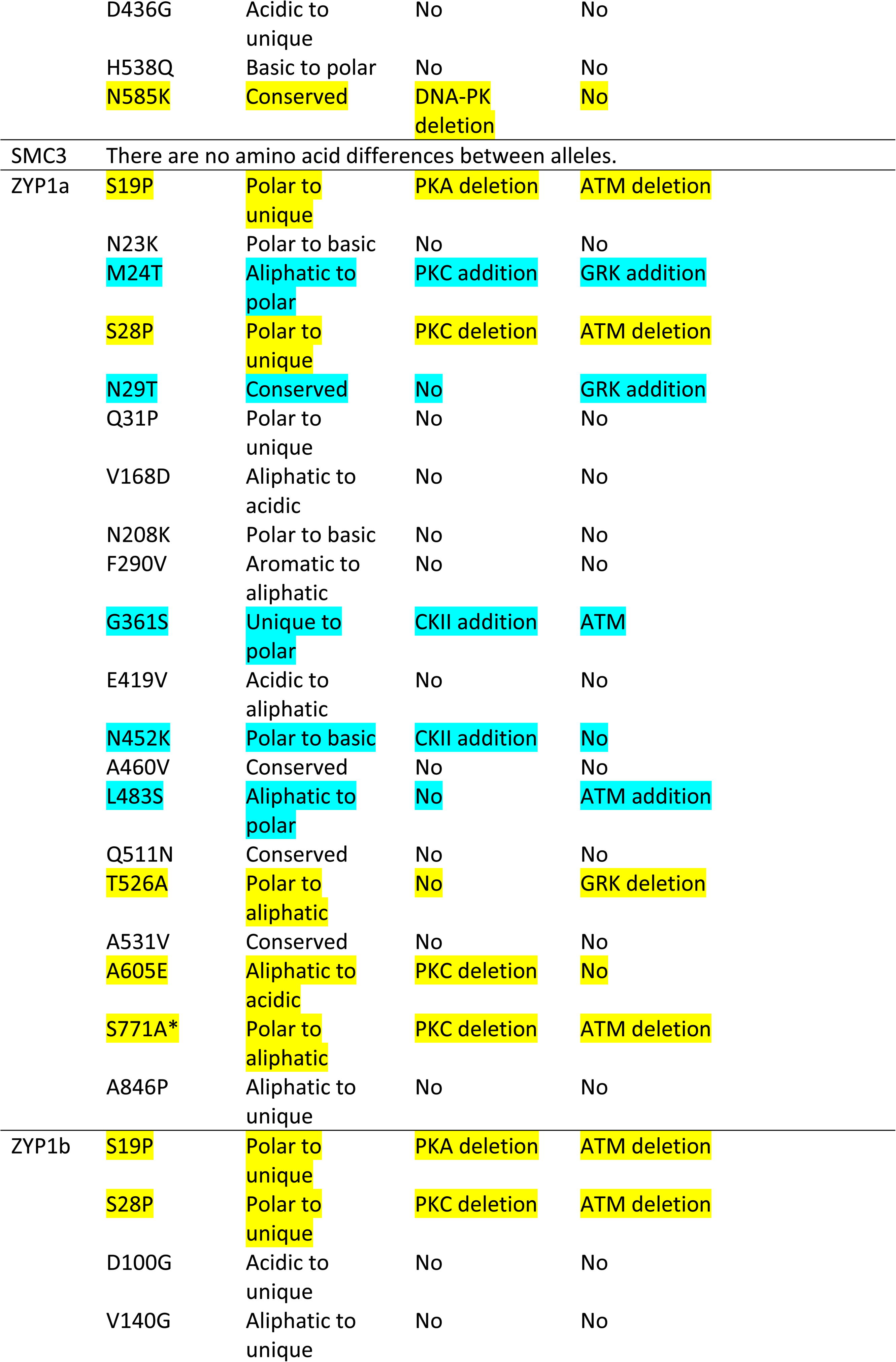
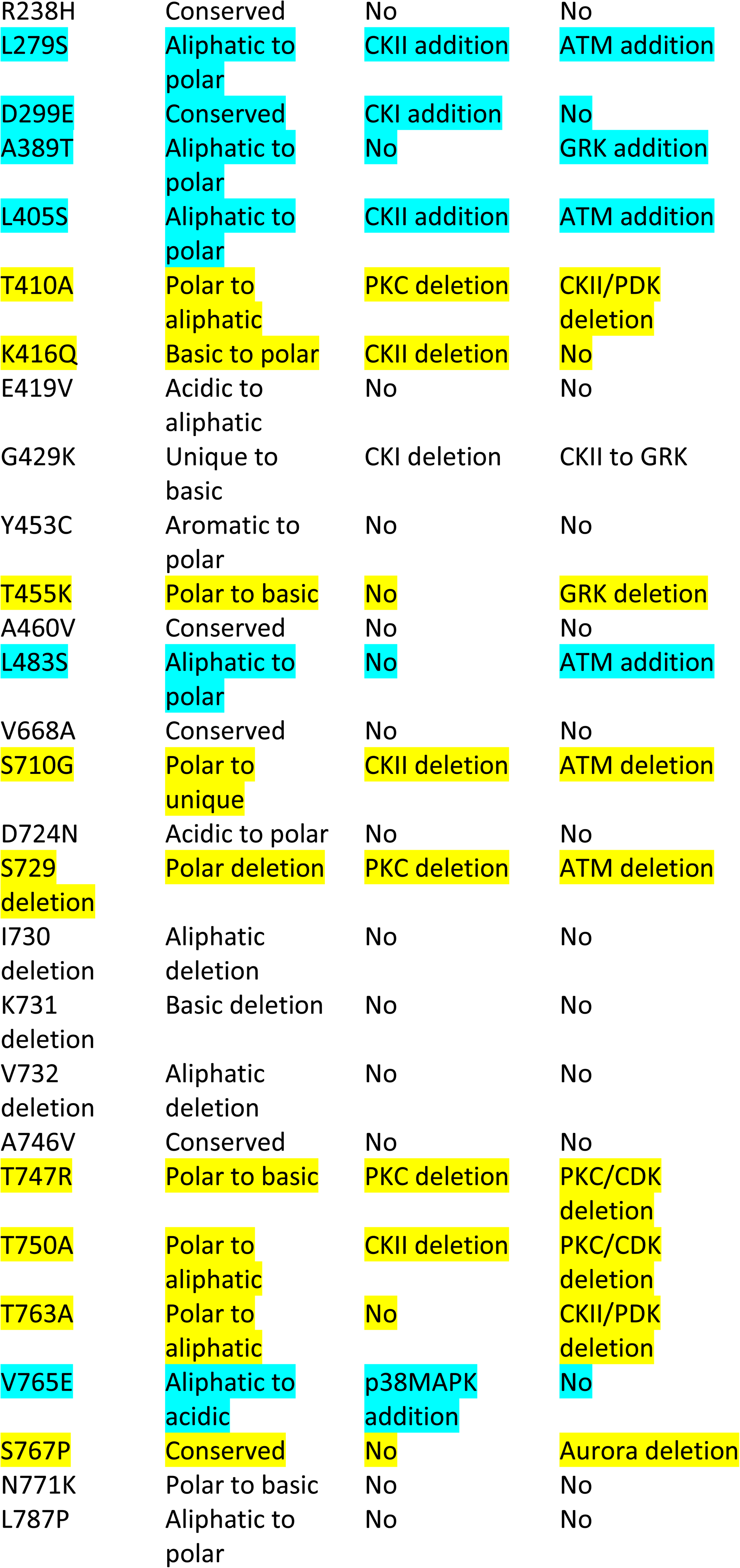
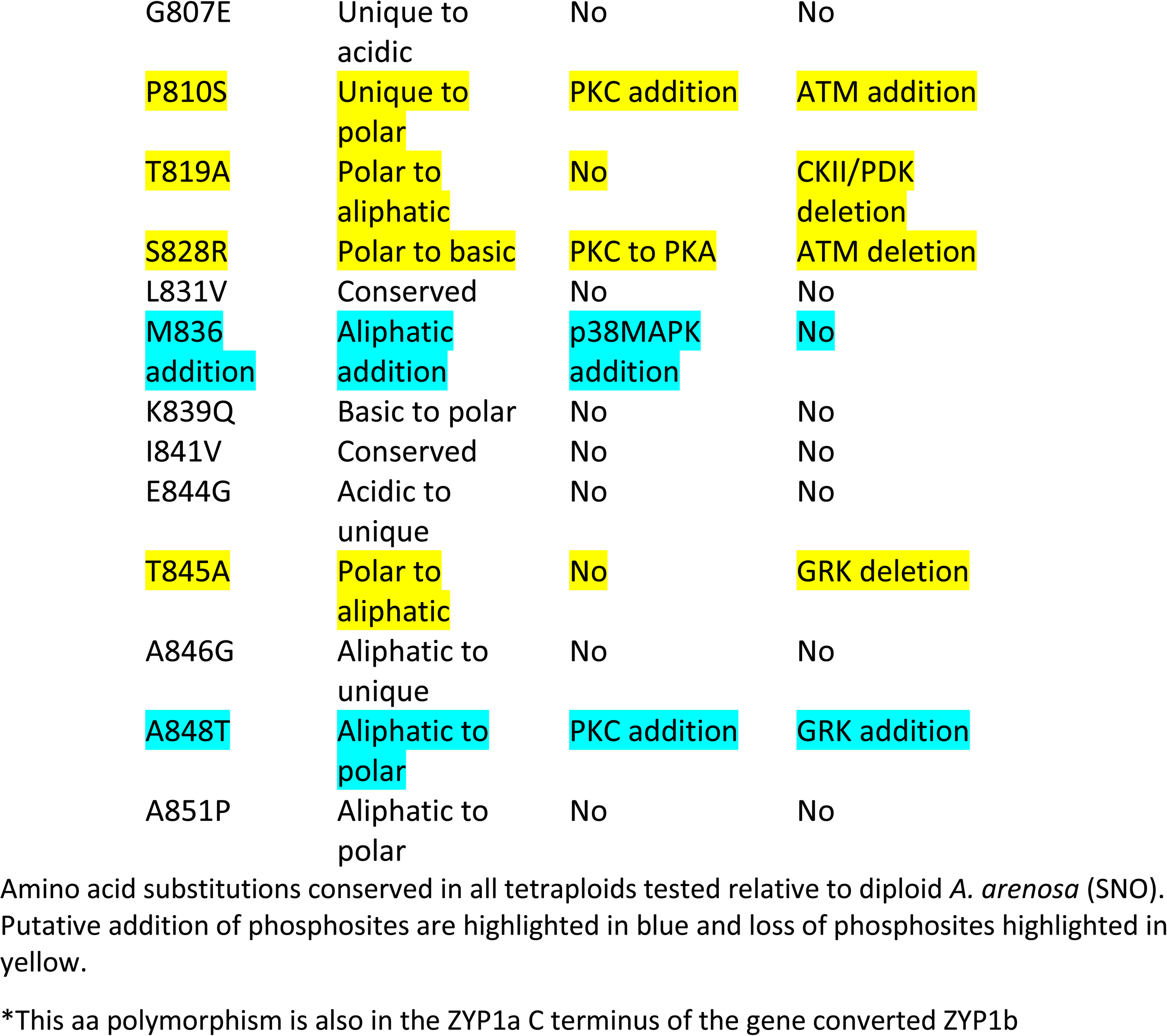

| Locus | Substitution | Property change | Phospho change (NetPhos 3.1) | Phospho change (KinasePhos2.0) | Domain |
| --- | --- | --- | --- | --- | --- |
| ASY1 | F273S | Aromatic to polar | No | Addition of ATM | PHD finger |
|  | R314H | Conserved | No | Increased probability of ATM in 4n | PHD finger |
|  | Q567R | Polar to basic | DNA-PK deletion | Decreased probability of ATM in 4n |  |
| PRD3 | L35 deletion | Aliphatic deletion | No | No |  |
|  | R36 deletion | Basic deletion | No | No |  |
|  | S37 deletion | Polar deletion | DNA-PK deletion | ATM deletion |  |
|  | Q38 deletion | Polar deletion | No | No |  |
|  | S108Y | Polar to aromatic | CDC2 deletion | ATM deletion |  |
|  | L128 deletion | Aliphatic deletion | No | No |  |
|  | A129S | Aliphatic to polar | ATM addition | ATM addition |  |
|  | T263M | Polar to aliphatic | p38MAPK deletion | CDC2 deletion |  |
|  | T272K | Polar to basic | PKC deletion | GRK deletion |  |
|  | V273A | Conserved | No | No |  |
|  | P309T | Unique to polar | No | GRK addition |  |
|  | Q310R | Polar to basic | No | No |  |
|  | R325S | Basic to polar | CDC2 addition | ATM addition |  |
|  | F375C | Aromatic to polar | No | No |  |
| REC8 | L224R | Aliphatic to basic | No | No |  |
|  | H225Q | Basic to polar | No | No |  |
|  | A256V | Conserved | No | No |  |
|  | A276T | Aliphatic to polar | PKC addition | GRK addition |  |
|  | S351T | Conserved | No | ATM to GRK |  |
|  | R353H | Conserved | No | No |  |
|  | Q370H | Polar to basic | No | No |  |
|  | Q423H | Polar to basic | No | No |  |

|  |  |  |  |  |
| --- | --- | --- | --- | --- |
|  | D436G | Acidic to unique | No | No |
|  | H538Q | Basic to polar | No | No |
|  | N585K | Conserved | DNA-PK deletion | No |
| SMC3 | There are no amino acid differences between alleles. |  |  |  |
| ZYP1a | S19P | Polar to unique | PKA deletion | ATM deletion |
|  | N23K | Polar to basic | No | No |
|  | M24T | Aliphatic to polar | PKC addition | GRK addition |
|  | S28P | Polar to unique | PKC deletion | ATM deletion |
|  | N29T | Conserved | No | GRK addition |
|  | Q31P | Polar to unique | No | No |
|  | V168D | Aliphatic to acidic | No | No |
|  | N208K | Polar to basic | No | No |
|  | F290V | Aromatic to aliphatic | No | No |
|  | G361S | Unique to polar | CKII addition | ATM |
|  | E419V | Acidic to aliphatic | No | No |
|  | N452K | Polar to basic | CKII addition | No |
|  | A460V | Conserved | No | No |
|  | L483S | Aliphatic to polar | No | ATM addition |
|  | Q511N | Conserved | No | No |
|  | T526A | Polar to aliphatic | No | GRK deletion |
|  | A531V | Conserved | No | No |
|  | A605E | Aliphatic to acidic | PKC deletion | No |
|  | S771A* | Polar to aliphatic | PKC deletion | ATM deletion |
|  | A846P | Aliphatic to unique | No | No |
| ZYP1b | S19P | Polar to unique | PKA deletion | ATM deletion |
|  | S28P | Polar to unique | PKC deletion | ATM deletion |
|  | D100G | Acidic to unique | No | No |
|  | V140G | Aliphatic to unique | No | No |

|  |  |  |  |
| --- | --- | --- | --- |
| R238H | Conserved | No | No |
| L279S | Aliphatic to polar | CKII addition | ATM addition |
| D299E | Conserved | CKI addition | No |
| A389T | Aliphatic to polar | No | GRK addition |
| L405S | Aliphatic to polar | CKII addition | ATM addition |
| T410A | Polar to aliphatic | PKC deletion | CKII/PDK deletion |
| K416Q | Basic to polar | CKII deletion | No |
| E419V | Acidic to aliphatic | No | No |
| G429K | Unique to basic | CKI deletion | CKII to GRK |
| Y453C | Aromatic to polar | No | No |
| T455K | Polar to basic | No | GRK deletion |
| A460V | Conserved | No | No |
| L483S | Aliphatic to polar | No | ATM addition |
| V668A | Conserved | No | No |
| S710G | Polar to unique | CKII deletion | ATM deletion |
| D724N | Acidic to polar | No | No |
| S729 deletion | Polar deletion | PKC deletion | ATM deletion |
| I730 deletion | Aliphatic deletion | No | No |
| K731 deletion | Basic deletion | No | No |
| V732 deletion | Aliphatic deletion | No | No |
| A746V | Conserved | No | No |
| T747R | Polar to basic | PKC deletion | PKC/CDK deletion |
| T750A | Polar to aliphatic | CKII deletion | PKC/CDK deletion |
| T763A | Polar to aliphatic | No | CKII/PDK deletion |
| V765E | Aliphatic to acidic | p38MAPK addition | No |
| S767P | Conserved | No | Aurora deletion |
| N771K | Polar to basic | No | No |
| L787P | Aliphatic to polar | No | No |
| G807E | Unique to acidic | No | No |
| P810S | Unique to polar | PKC addition | ATM addition |
| T819A | Polar to aliphatic | No | CKII/PDK deletion |
| S828R | Polar to basic | PKC to PKA | ATM deletion |
| L831V | Conserved | No | No |
| M836 addition | Aliphatic addition | p38MAPK addition | No |
| K839Q | Basic to polar | No | No |
| I841V | Conserved | No | No |
| E844G | Acidic to unique | No | No |
| T845A | Polar to aliphatic | No | GRK deletion |
| A846G | Aliphatic to unique | No | No |
| A848T | Aliphatic to polar | PKC addition | GRK addition |
| A851P | Aliphatic to polar | No | No |

Amino acid substitutions conserved in all tetraploids tested relative to diploid *A. arenosa* (SNO). Putative addition of phosphosites are highlighted in blue and loss of phosphosites highlighted in yellow.
\*This aa polymorphism is also in the ZYP1a C terminus of the gene converted ZYP1b

**S5 Table.**
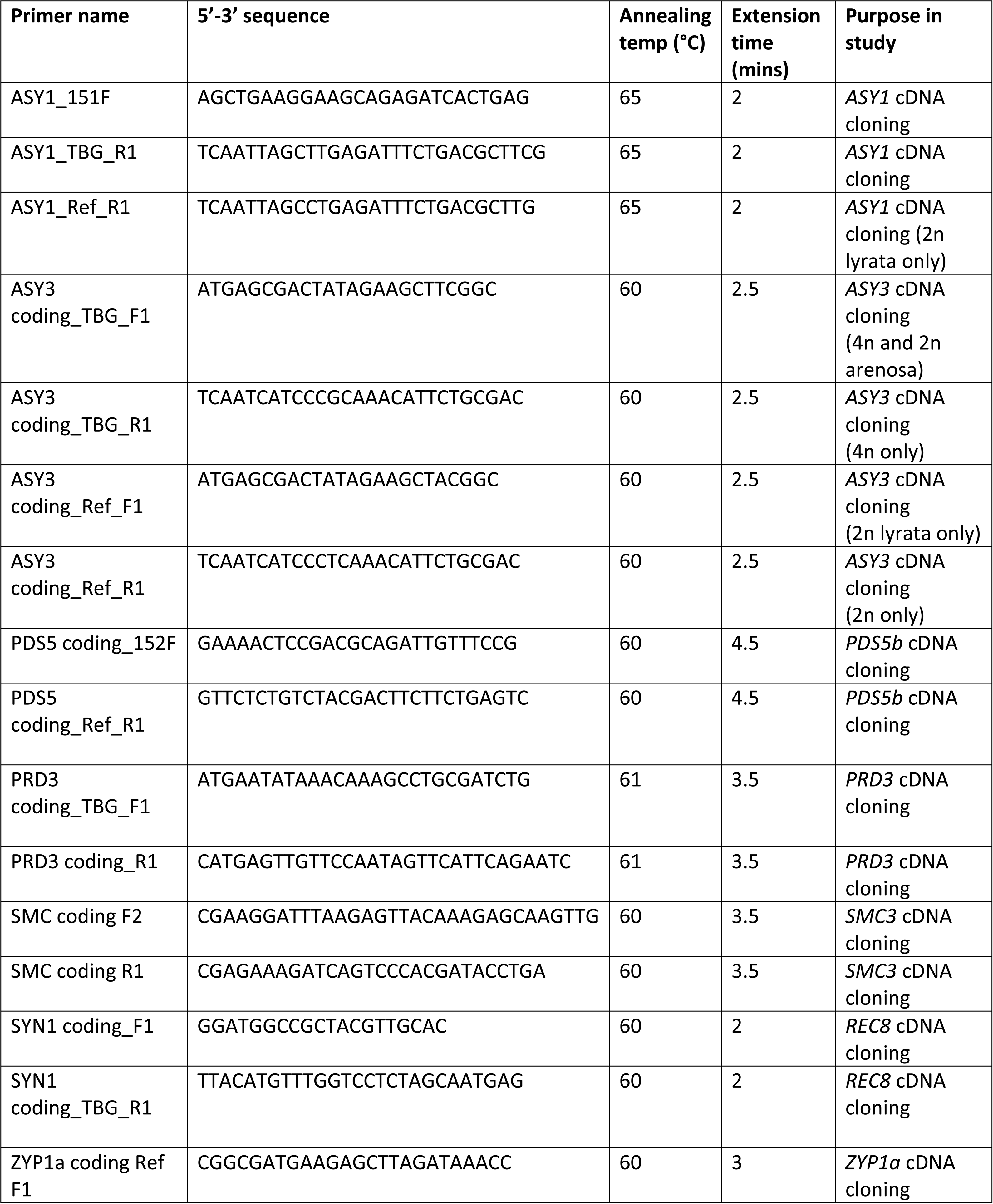
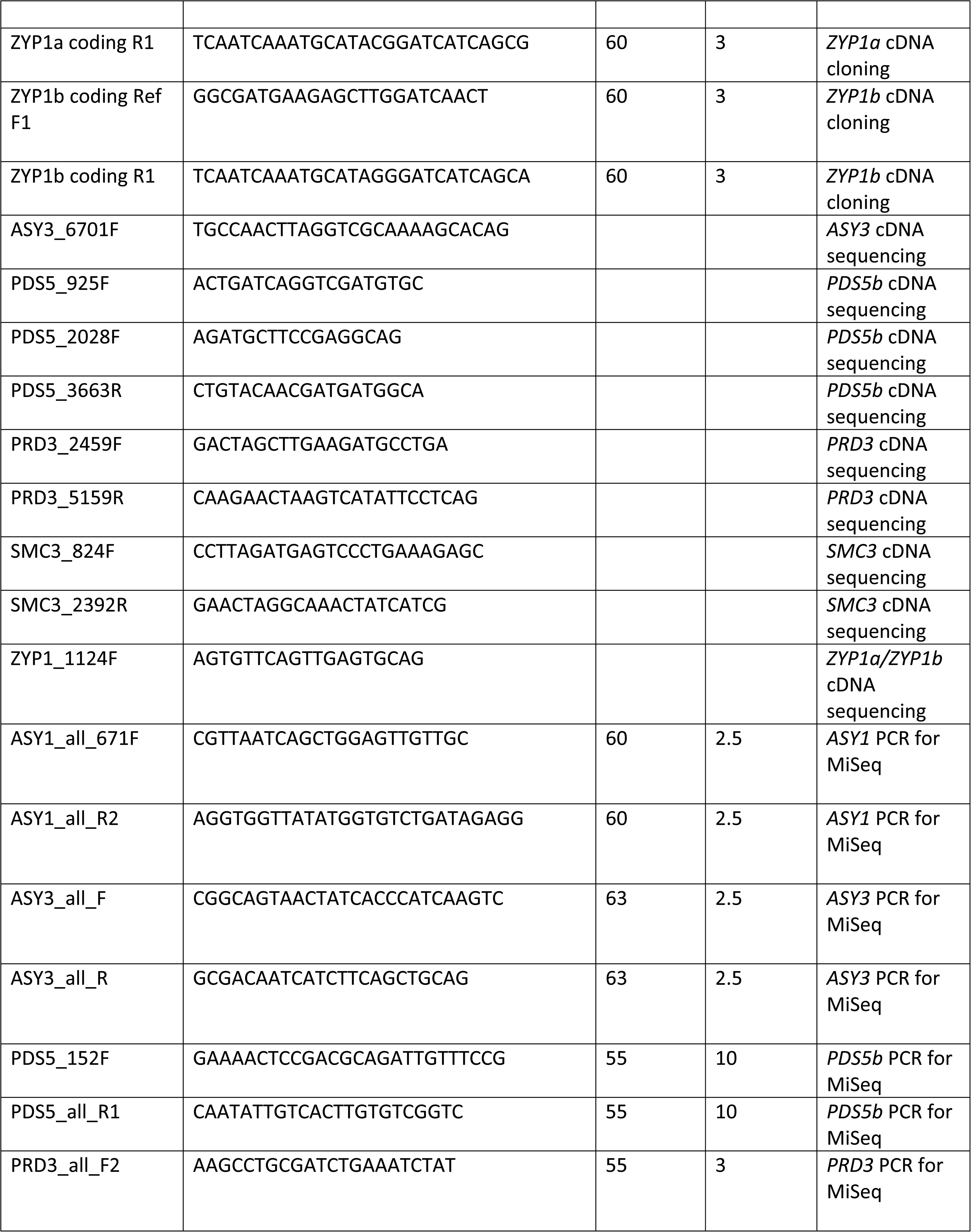
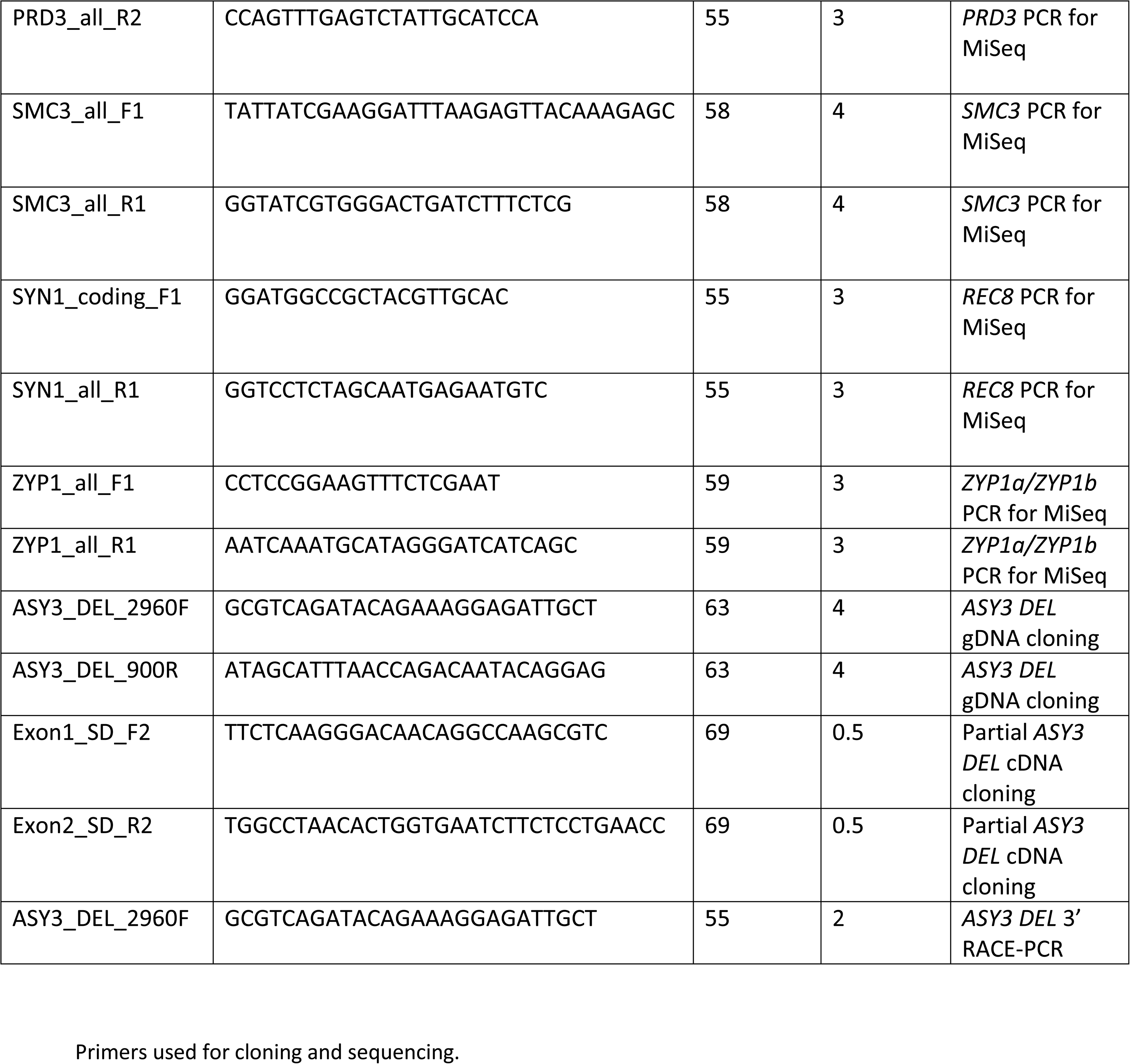

**S6 Table.**

| **EnsemblPlants ID** | **Gene** | **Location** |
| --- | --- | --- |
| Scaffold_201546.1 | *ASY1* | Chr2:12427364:12431683 |
| Fgenesh2_kg.4_2929 | *ASY3* | Chr4:22845898:22850613 |
| Scaffold_202722.1 | *PDS5b* | Chr2:17706052:17715900 |
| Scaffold_0001_57 | *PRD3* | Chr1:220193:225616 |
| Fgenesh1_pg.C_scaffold_4000806 | *SMC3* | Chr4:11123922:11131646 |
| Fgenesh1_pm.C_scaffold_6000400 | *REC8* | Chr6:2001440:2005979 |
| Fgenesh00000004233 | *ZYP1a* | Chr1:9779047:9784303 |
| Scaffold_102525.1 | *ZYP1b* | Chr1:9786597:9791542 |

Meiosis genes from *A.lyrata* reference genome

**S7 Table.**
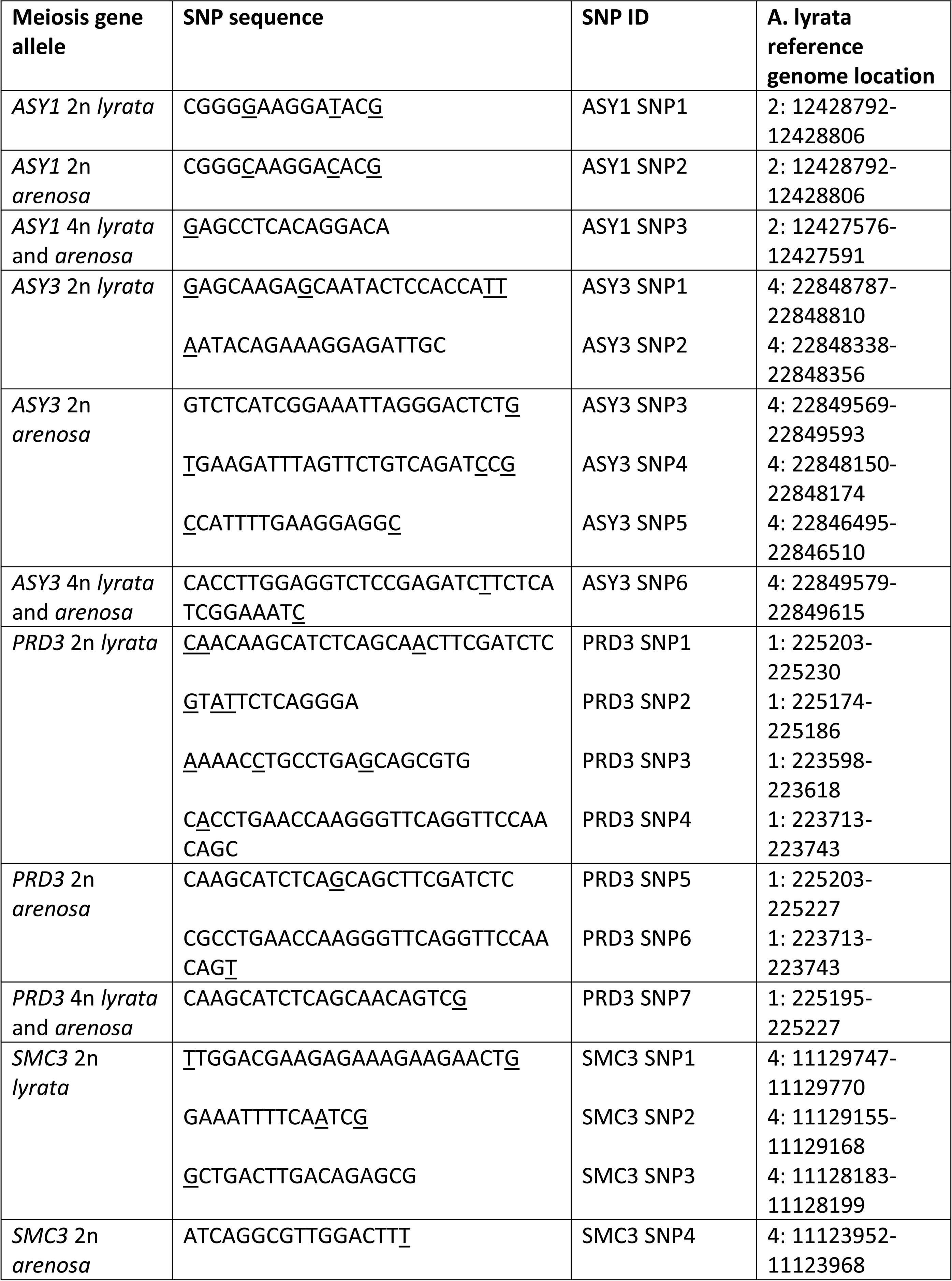
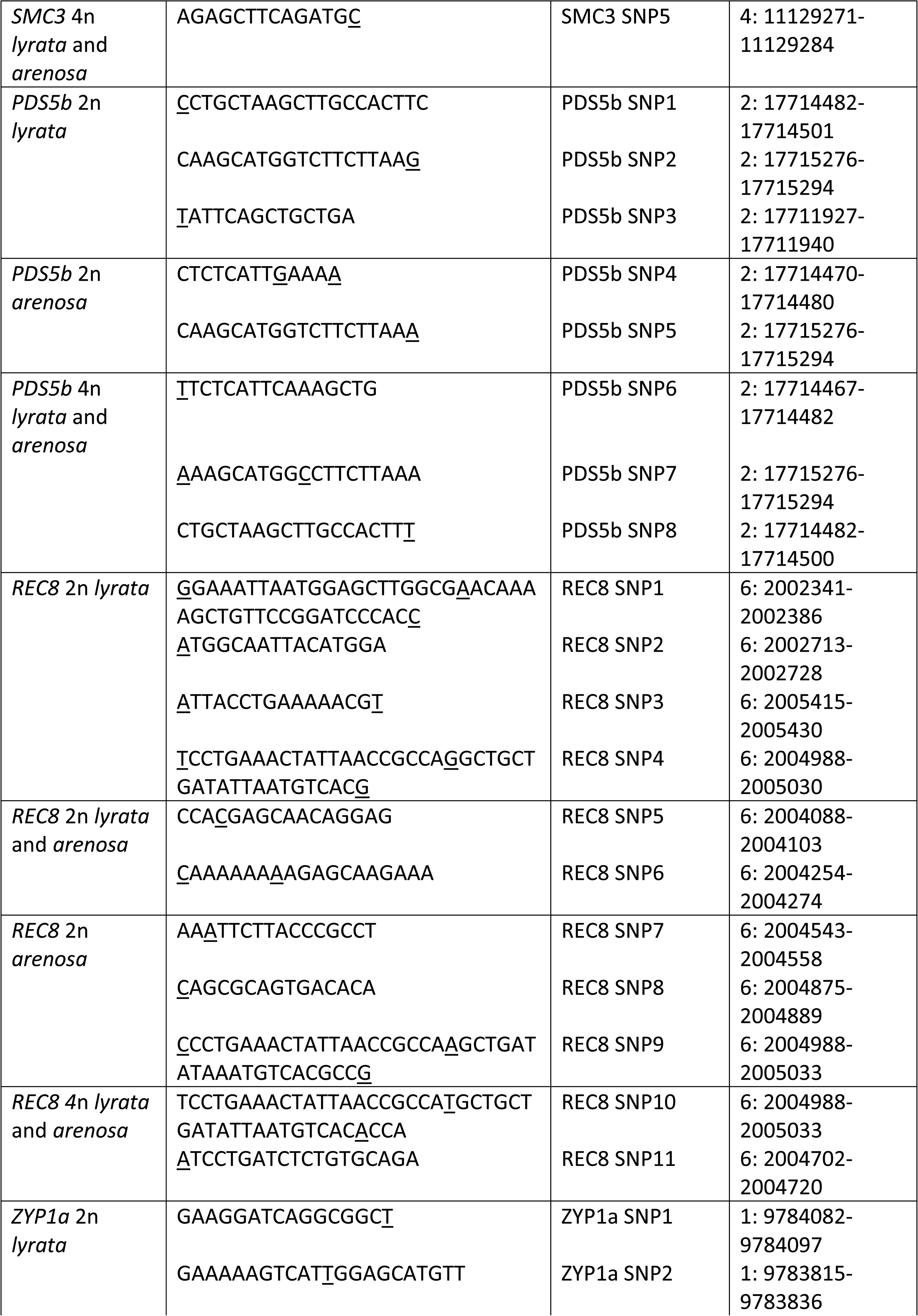
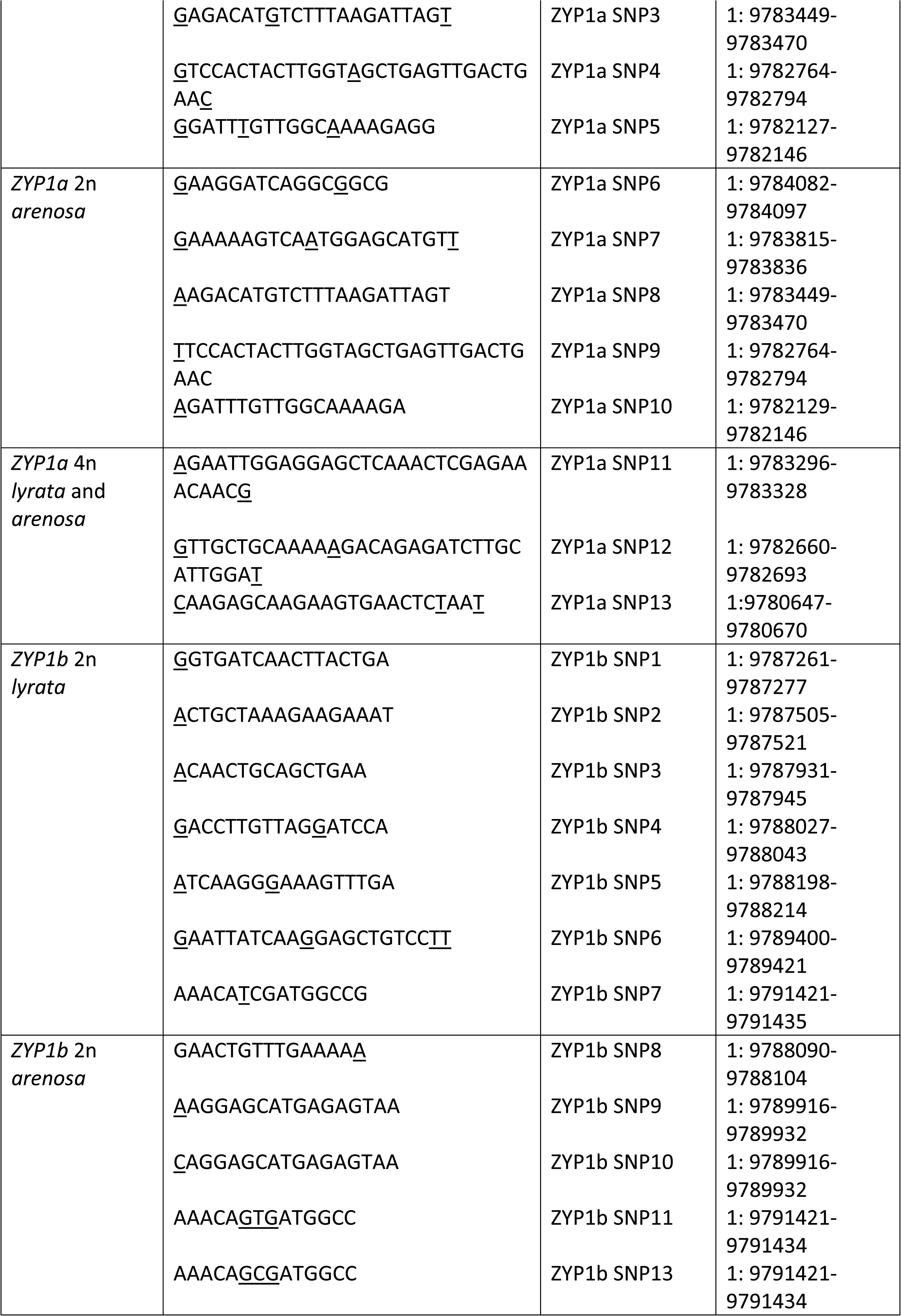
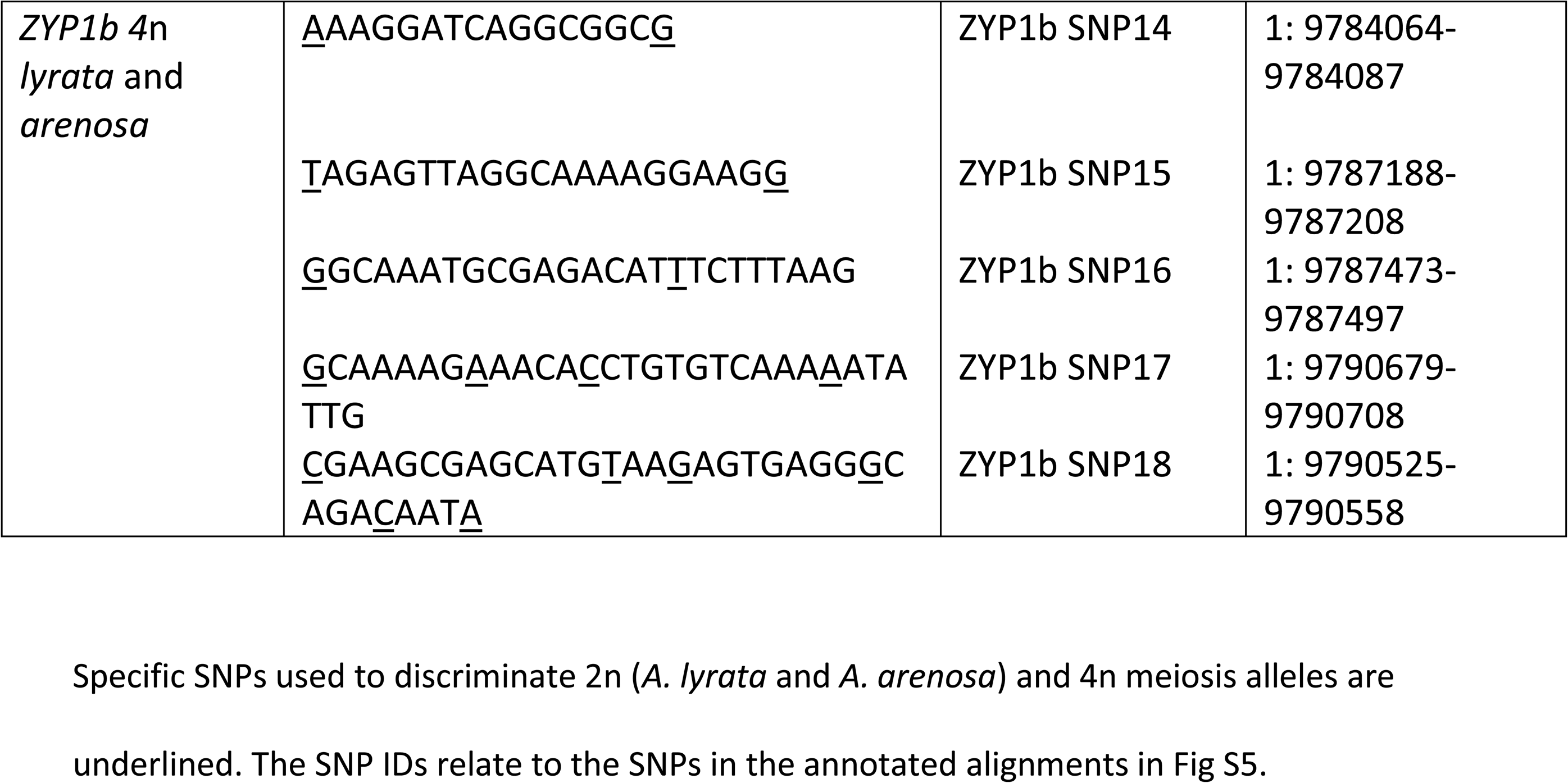

---

**S1 Fig.**
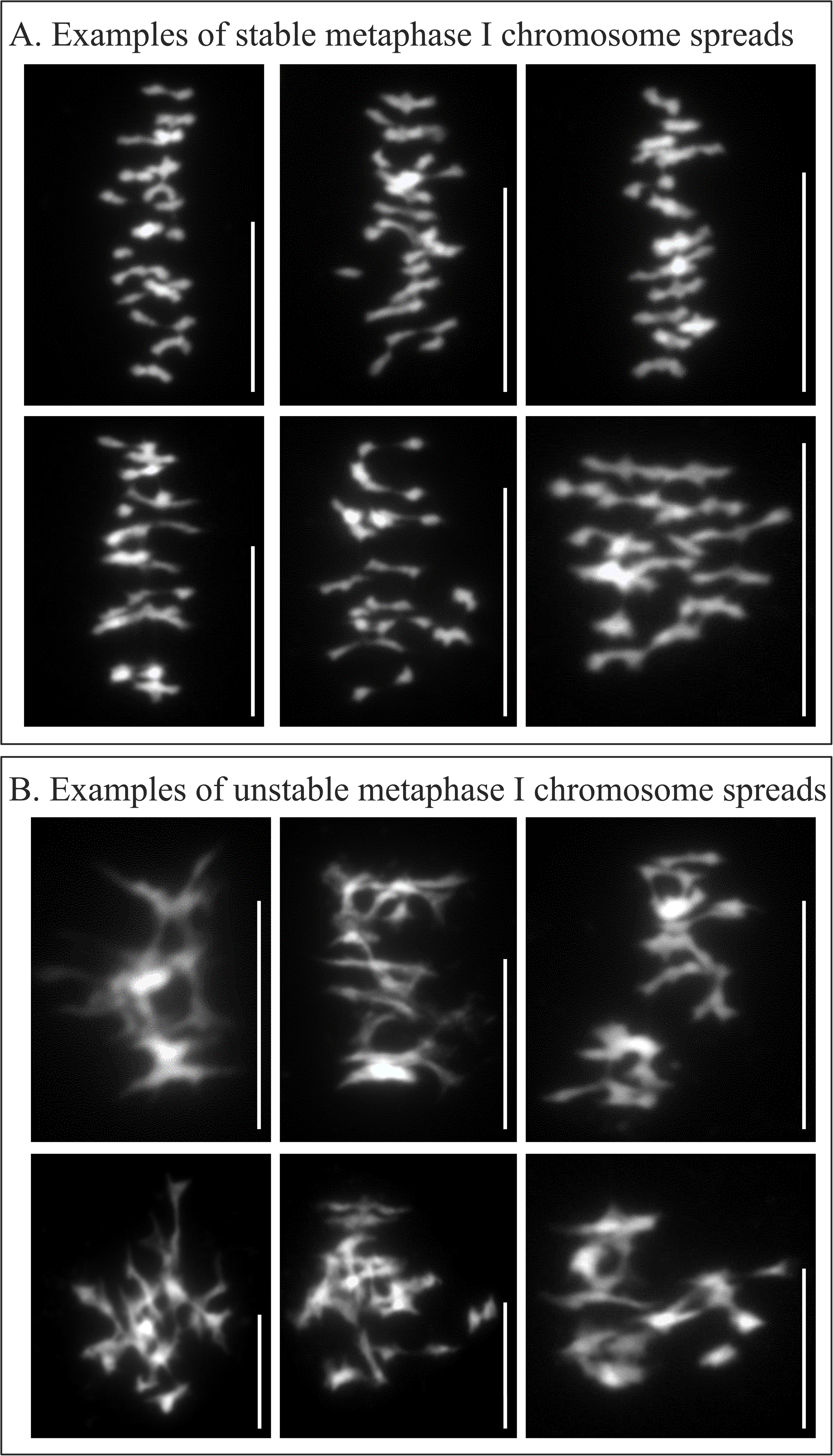
Examples of stable (A) and unstable (B) male meiotic metaphase 1 chromosome spreads. Scale bar = 10µM

**SFig 2.**
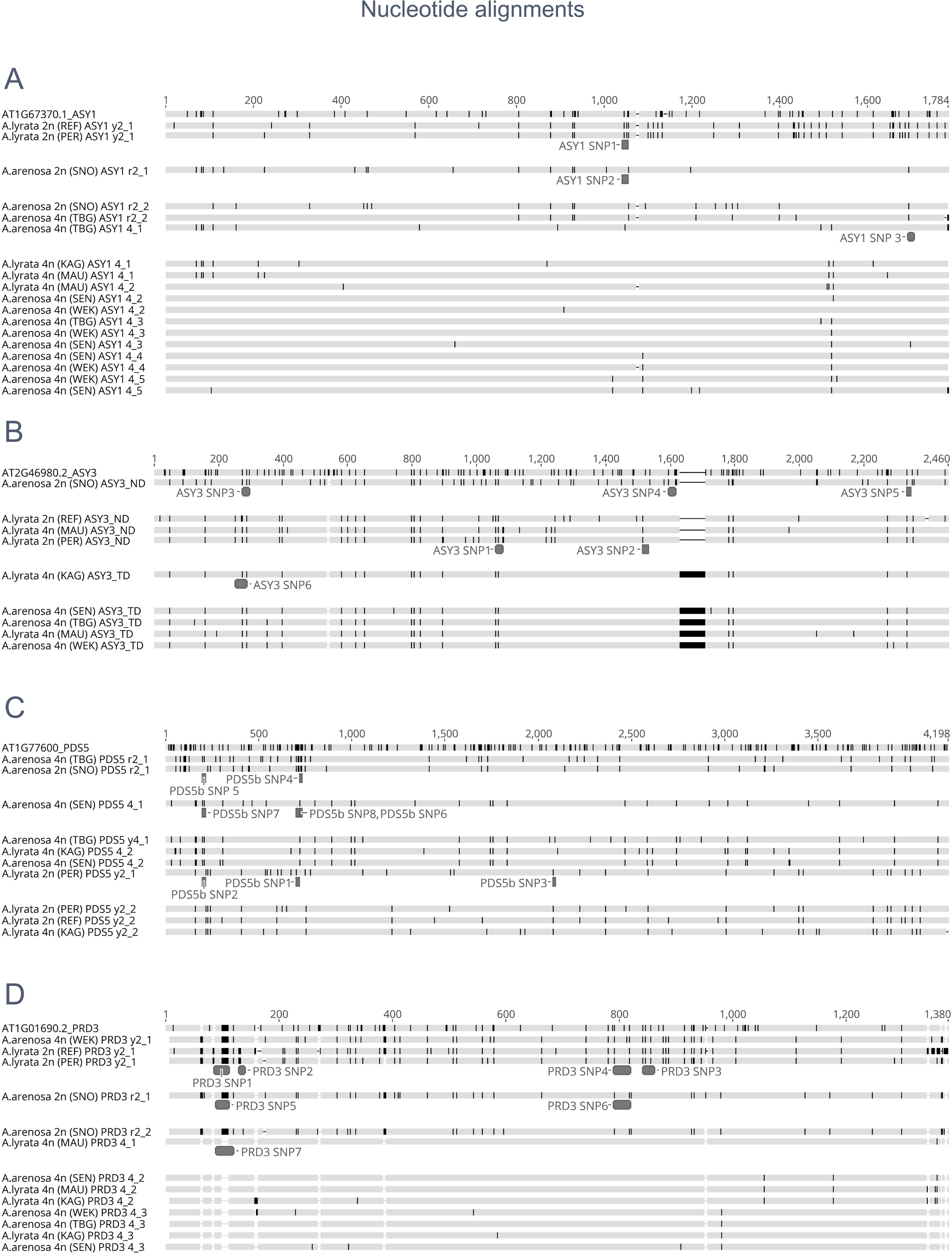
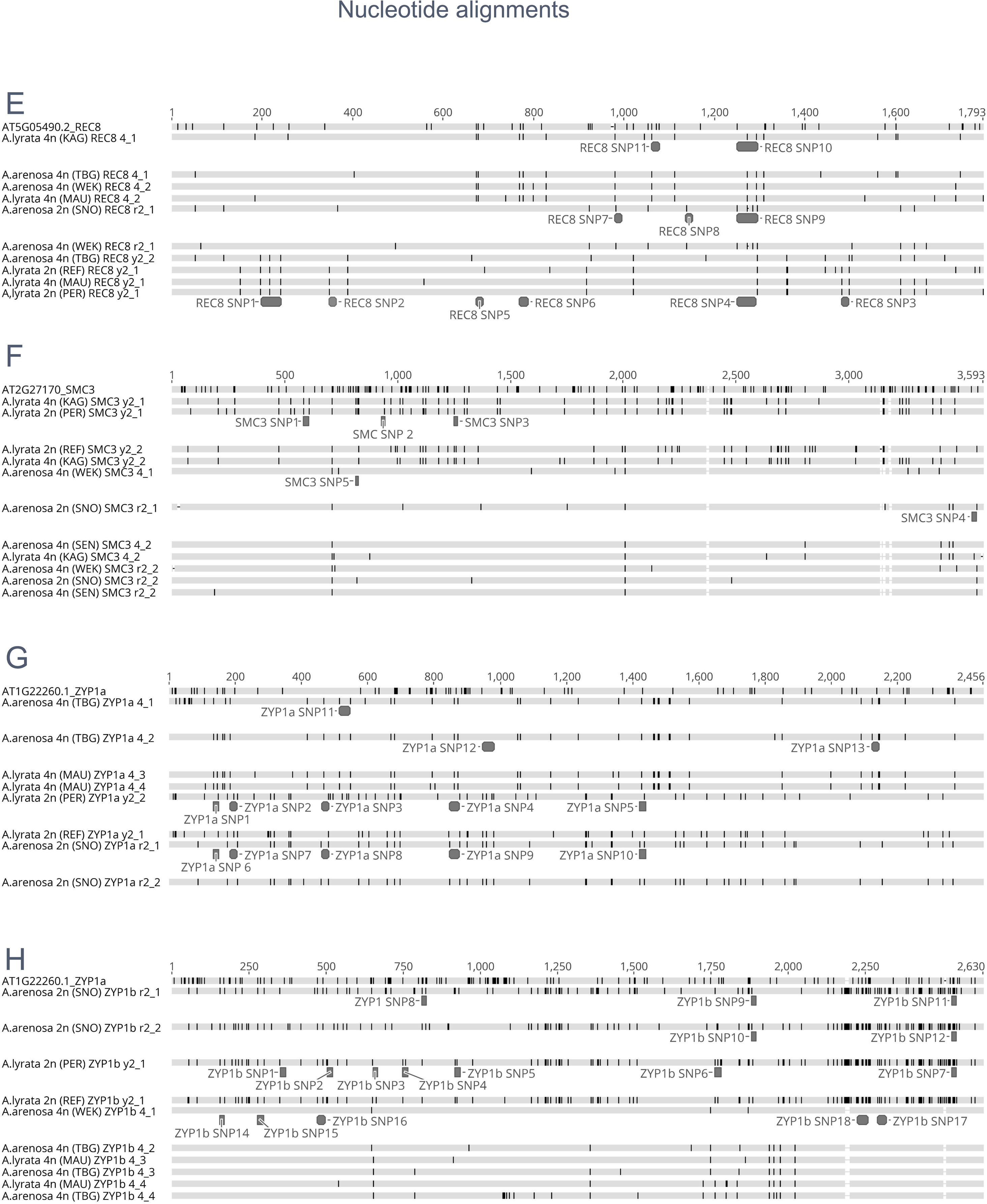
Nucleotide alignments of *ASY1* (A), *ASY3* (B), *PDS5b* (C), *PRD3* (D), *REC8* (E), *SMC3* (F), *ZYP1a* (G) and *ZYP1b* (H) in diploid and autotetraploid *A. arenosa* and *A. lyrata*, in addition to *A. thaliana*. The location of diploid and autotetraploid allele specific SNPs are indicated. Black bars indicate SNPs relative to the consensus sequence (not shown).

**SFig 3.**
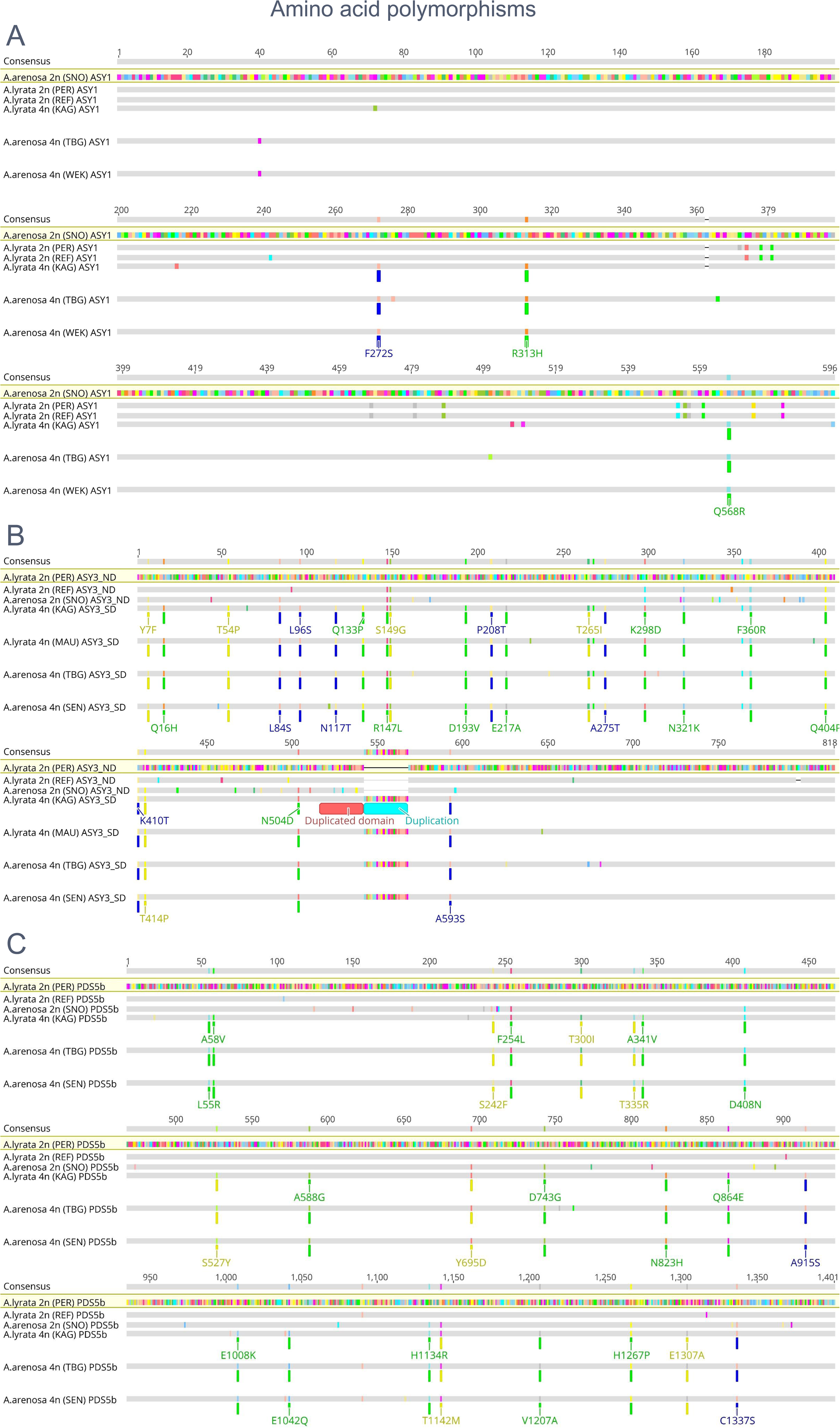
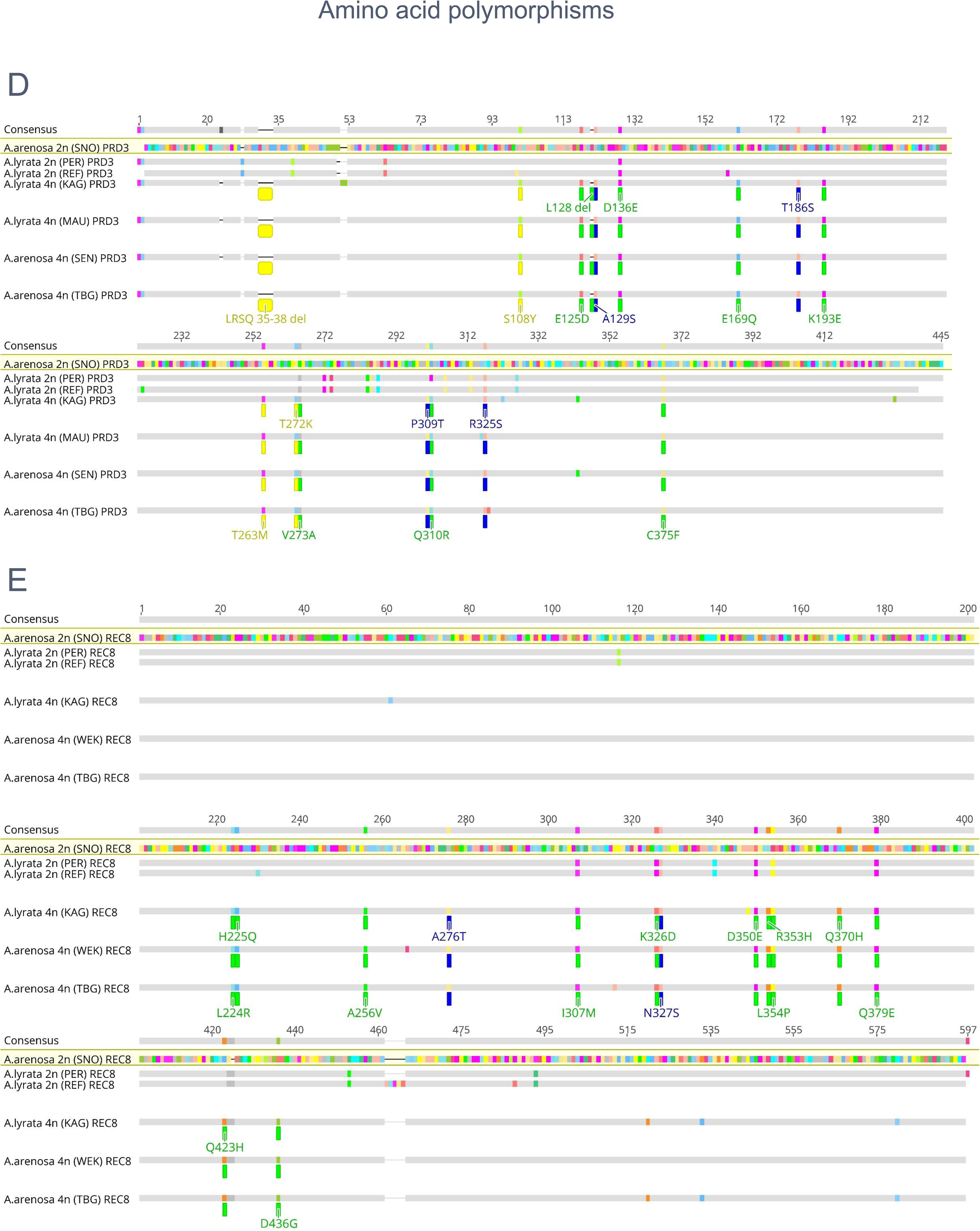
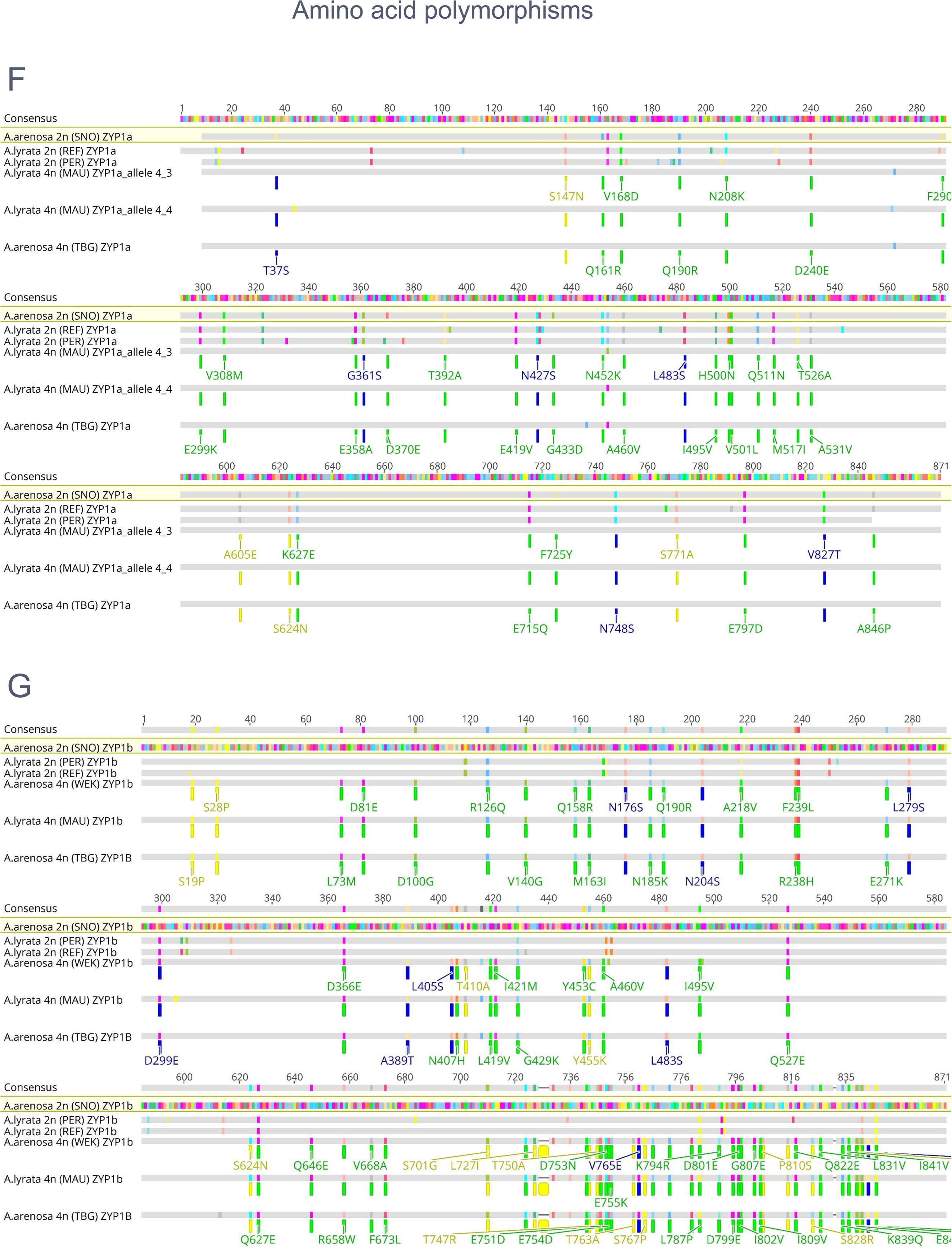
*In silico* translation alignments of ASY1 (A), ASY3 (B), PRD3 (C), PDS5b (D), REC8 (E), ZYP1a (F) and ZYP1b (G) showing conserved amino acid polymorphisms in autotetraploids compared to ancestral diploid alleles. Gains, losses and no change of predicted phosphosites are indicated in blue, yellow and green respectively.

**SFig 4.**
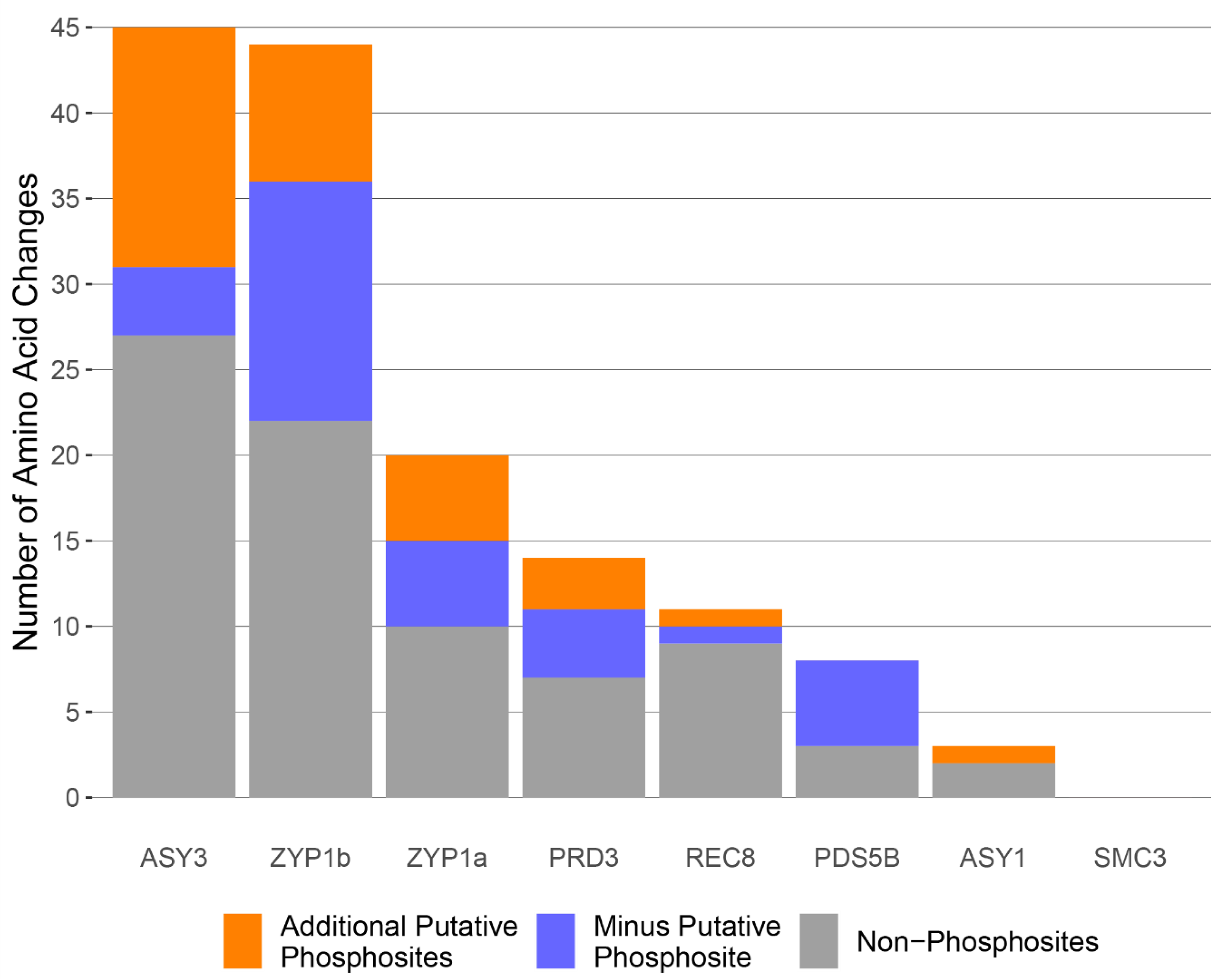
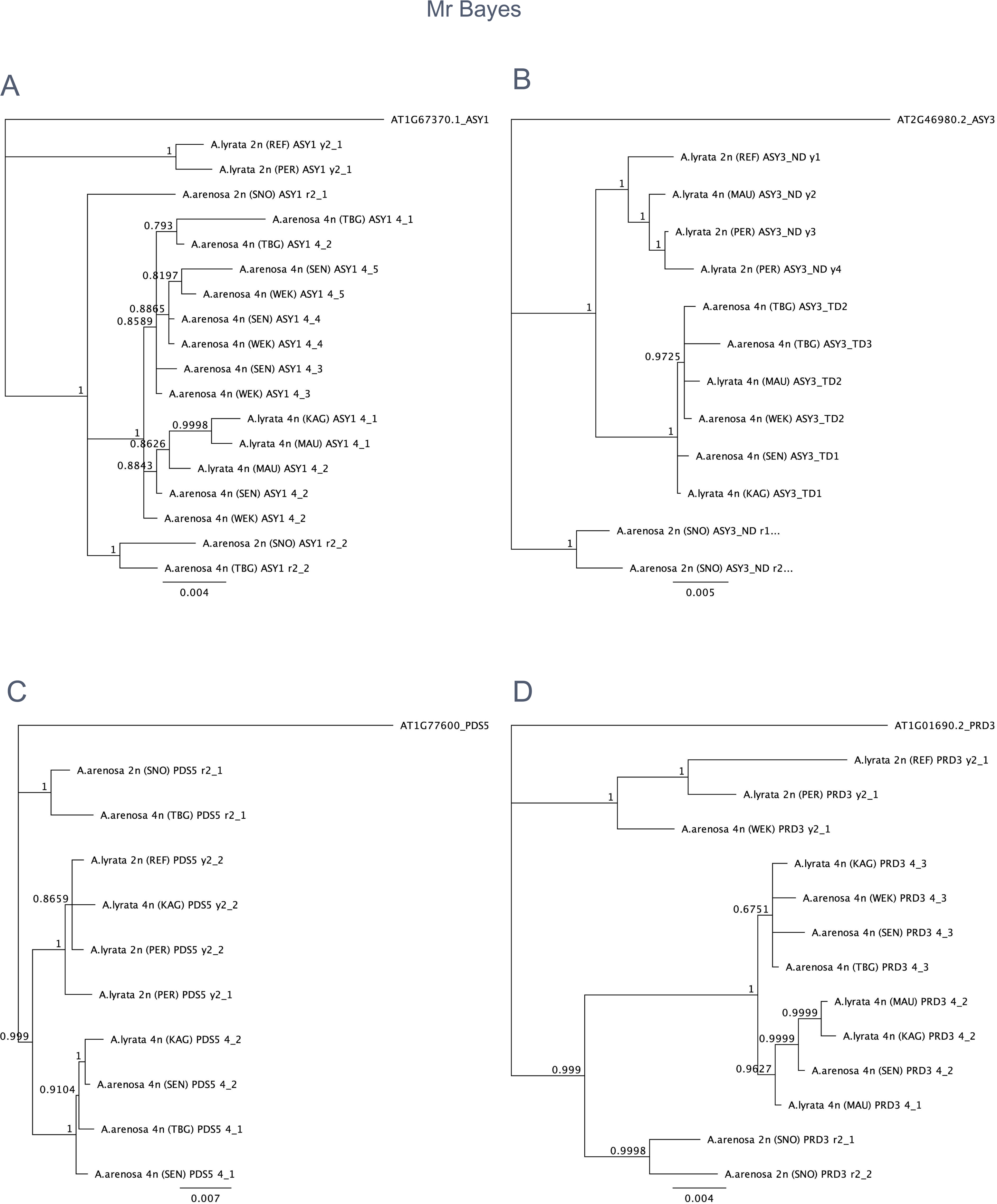
Summary of conserved amino acid polymorphisms in derived autotetraploid proteins compared to ancestral diploids including gains and losses of predicted serine/threonine phosphosites by KinasePhos2.0 and NetPhos3.1

**SFig 5.**
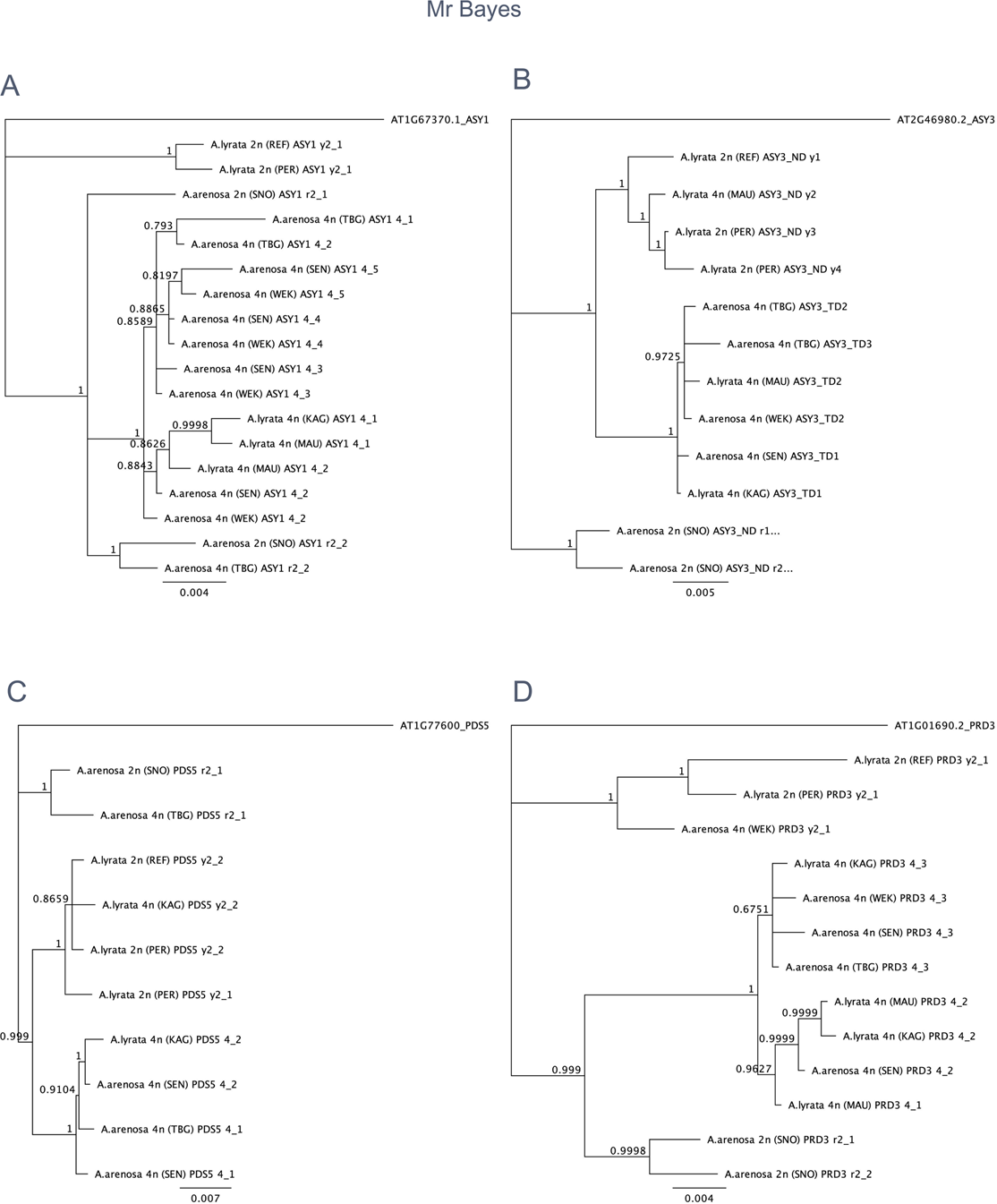
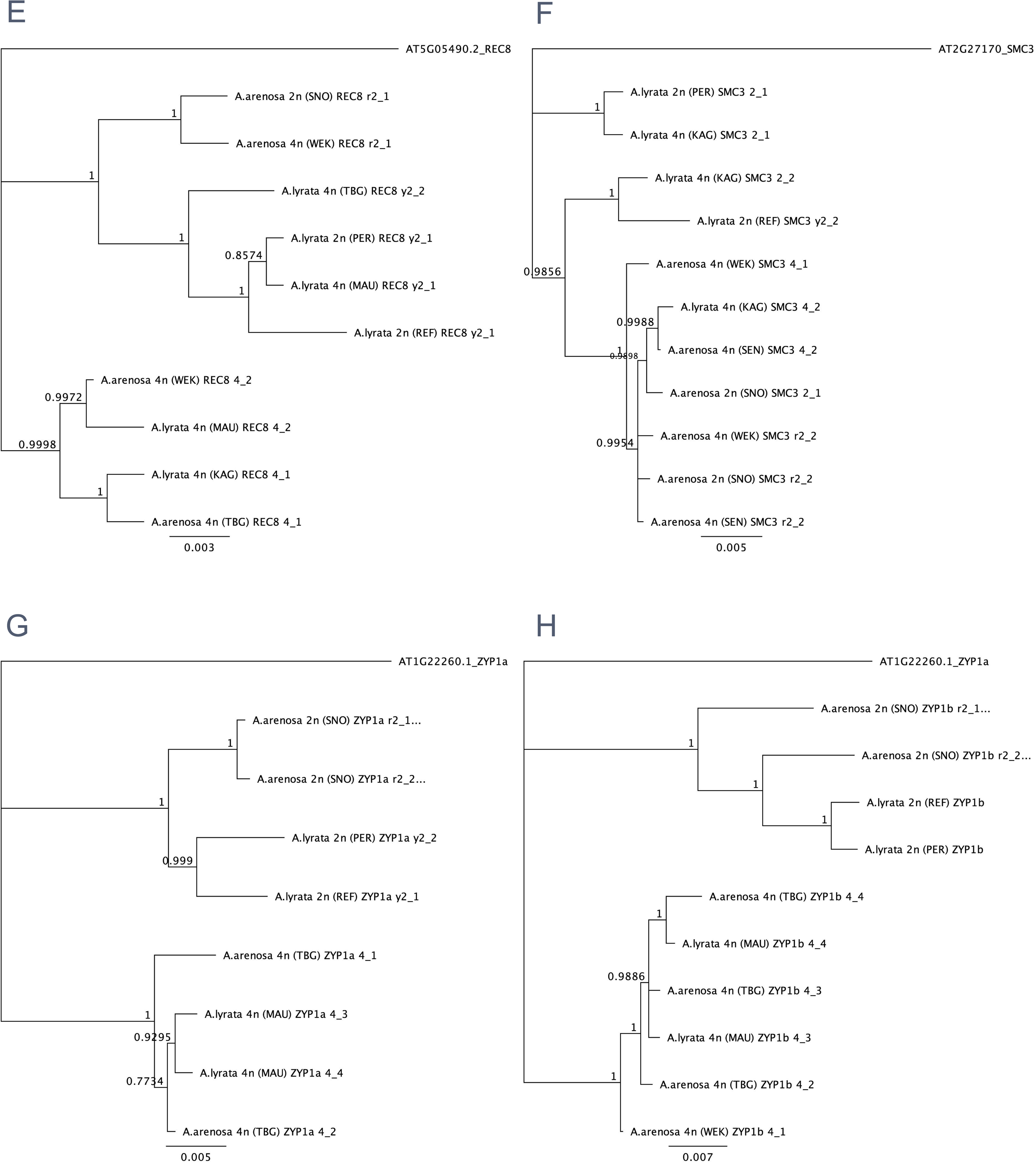
Phylogenetic trees of meiosis genes indicating origin of alleles selected in the autotetraploids constructed using a Bayesian approach with MrBayes v3.2.6. (A) *ASY1*, (B) *ASY3*, (C) *PDS5b*, (D) *PRD3,* (E) *REC8*, (F) *SMC3*, (G) *ZYP1a*, (H) *ZYP1b*. Bayesian posterior probabilities are indicated at the internodes of each branch (1000 replicates). The dissimilarity scale showing substitutions per nucleotide is located at the bottom of each tree.

**SFig 6.**
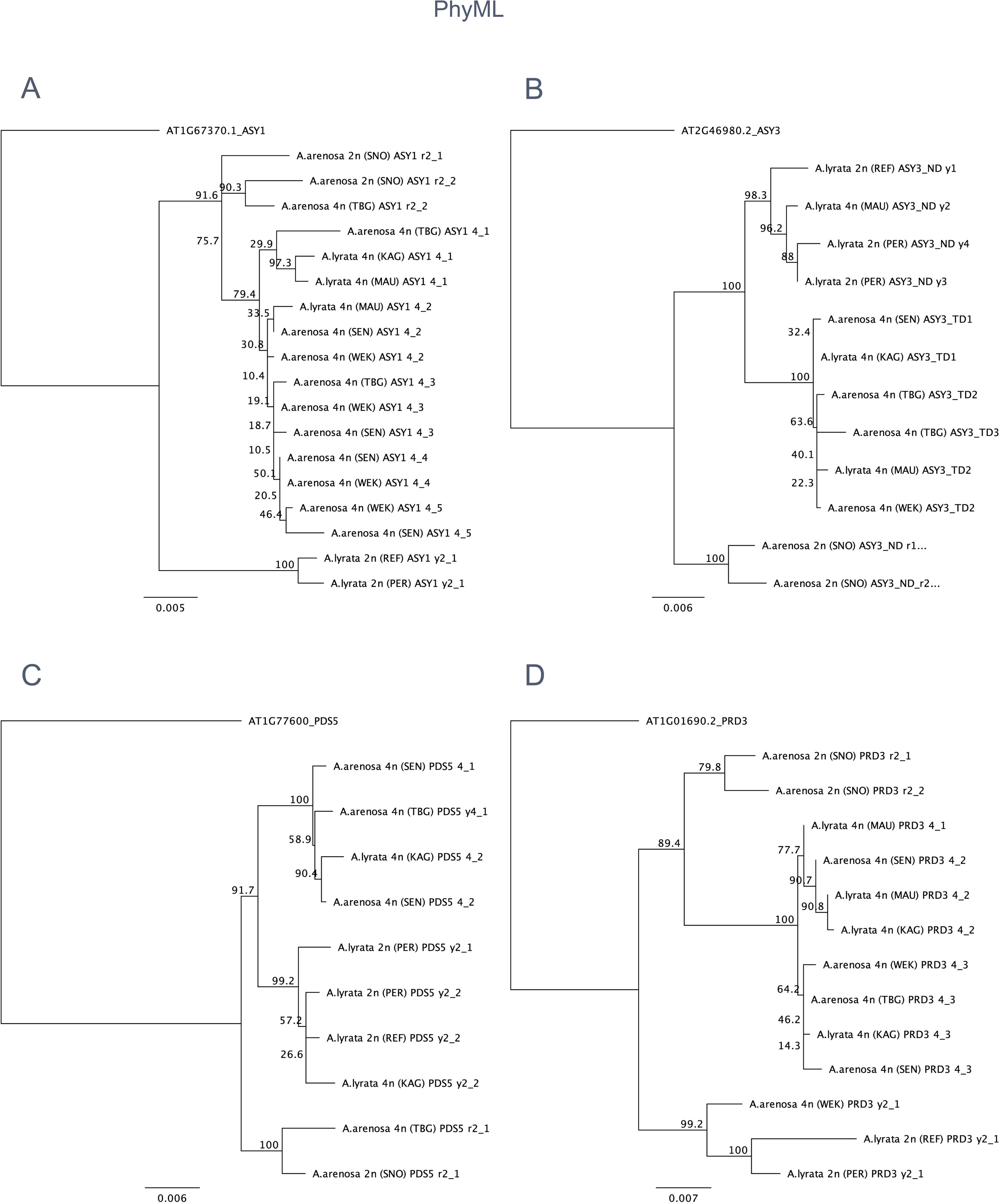
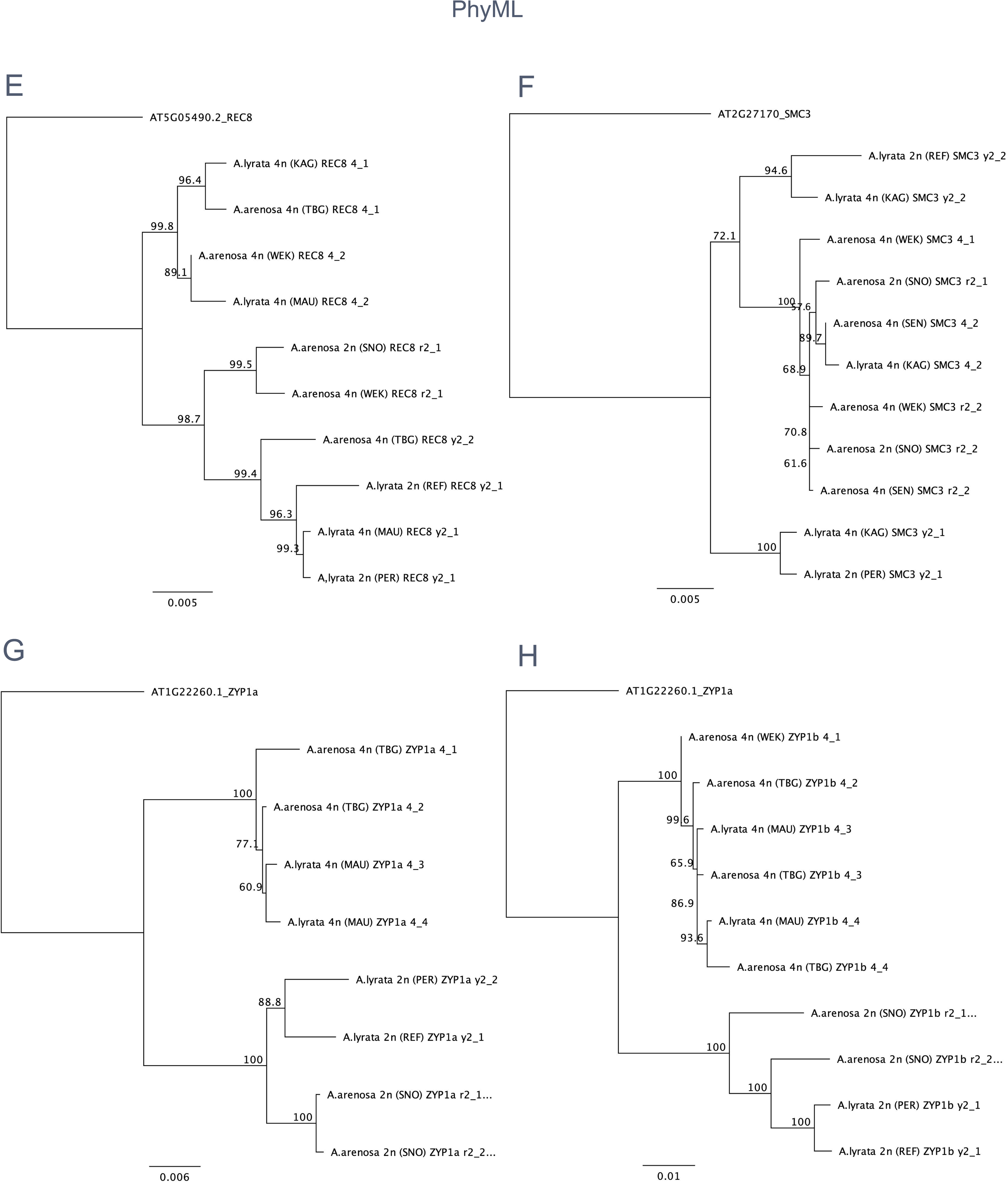
Phylogenetic trees of meiosis genes indicating origin of alleles selected in the autotetraploids constructed using a maximum likelihood approach with PhyML v3.3. (A) *ASY1*, (B) *ASY3*, (C) *PDS5b*, (D) *PRD3,* (E) *REC8*, (F) *SMC3*, (G) *ZYP1a*, (H) *ZYP1b*. Maximum likelihood bootstrap values are indicated at the internodes of each branch (1000 replicates). The dissimilarity scale showing substitutions per nucleotide is located at the bottom of each tree.

**SFig 7.**
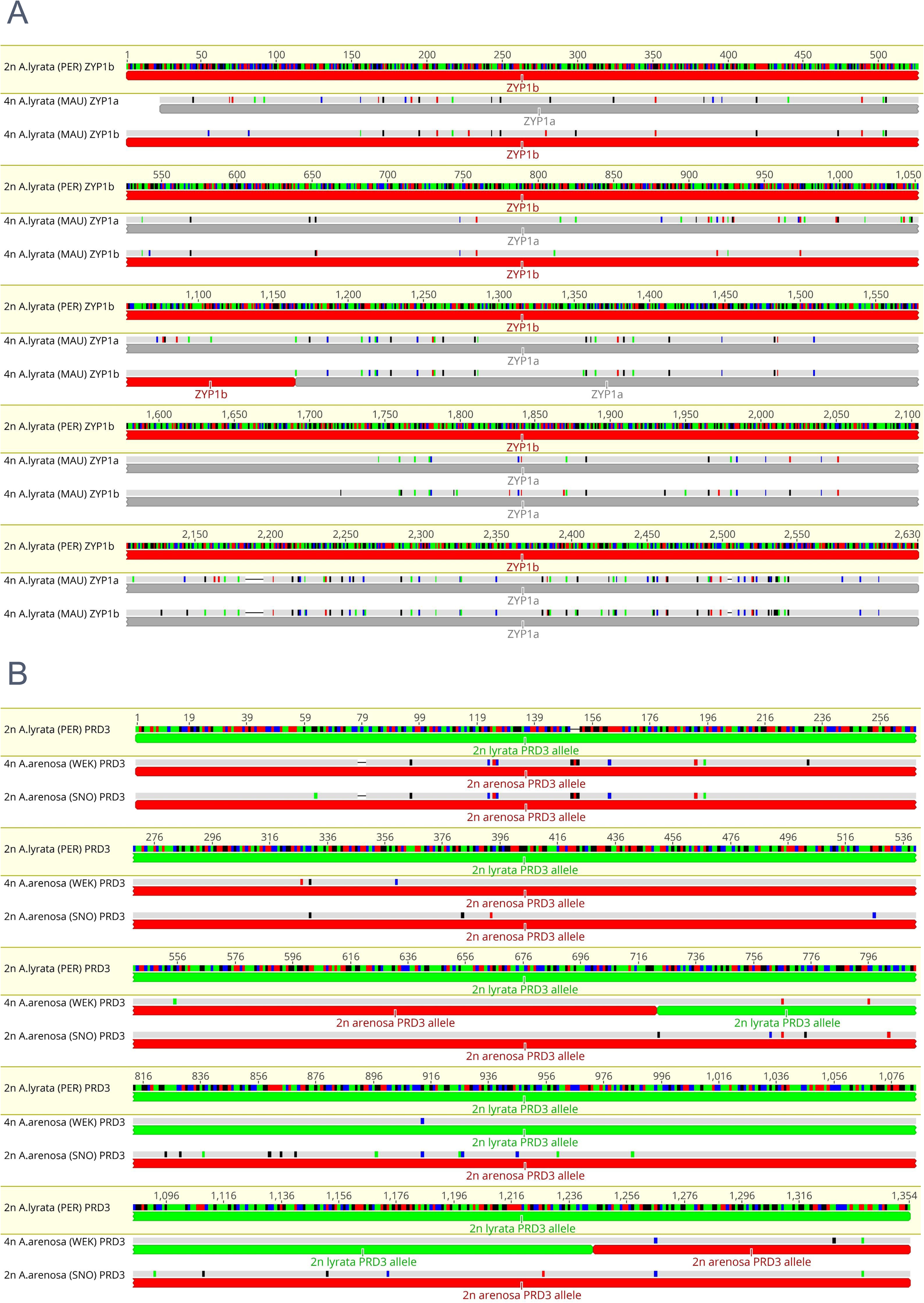
Nucleotide alignments showing examples of putative SC gene conversion-mediated protein polymorphisms. (A) *ZYP1* gene conversion (or CO) between *ZYP1a* (grey) and *ZYP1b* (red) in autotetraploid *ZYP1b* alleles, (B) gene conversion between PRD3 diploid *A. lyrata* (green) and *A. arenosa* (green) in tetraploid *A. arenosa*. Coloured bars in each sequence represent base specific SNPs relative to ancestral diploid alleles (Green = A, Blue = C, Black = G, Red = T).

**SFig 8.**
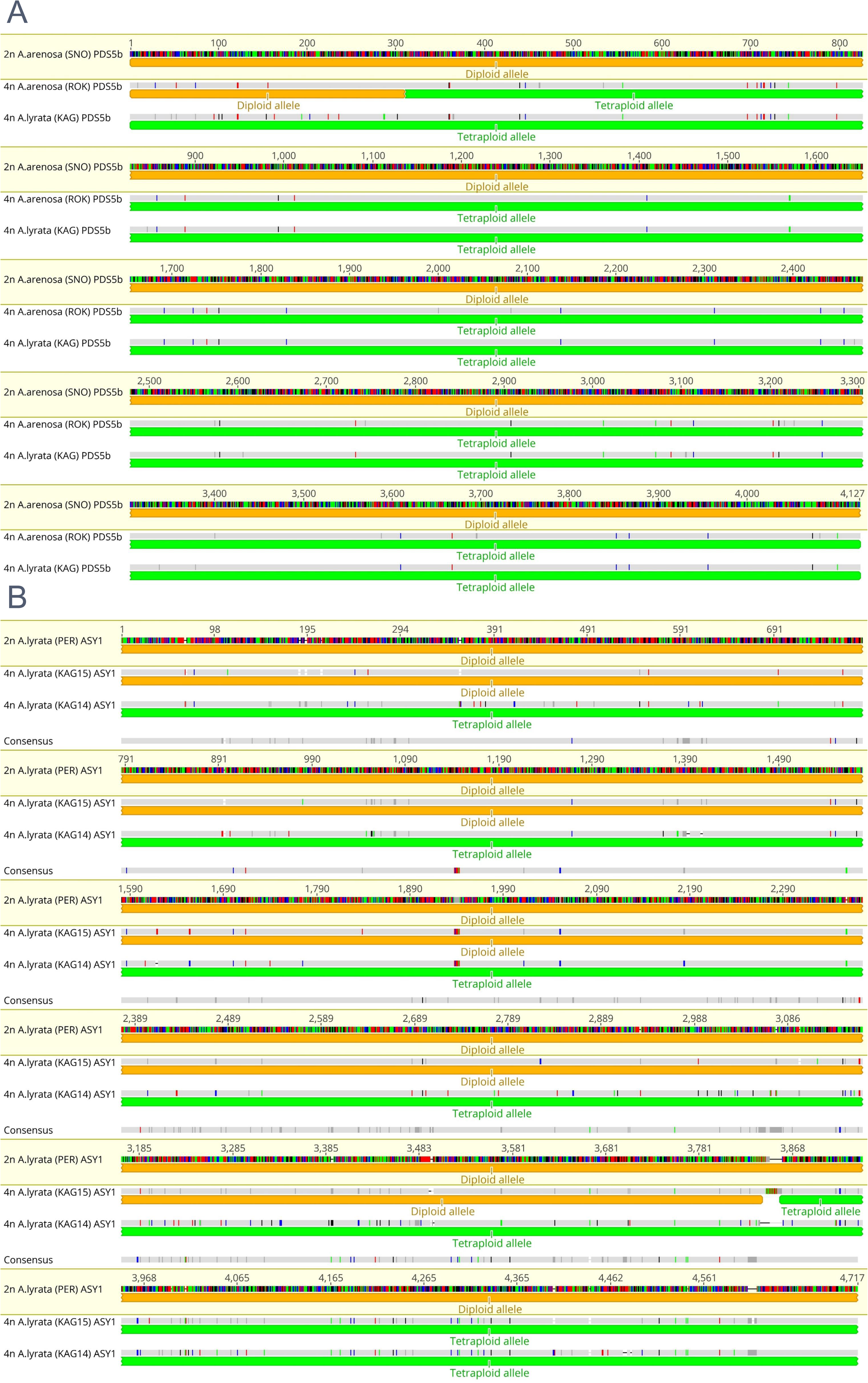
Nucleotide alignments showing examples of putative SC gene conversion-mediated protein polymorphisms. (A) gene conversion (or CO) between *PDS5b* diploid (yellow) and autotetraploid (green) alleles. (B) Gene conversion (or CO) between *ASY1* diploid *A. lyrata* (yellow) and autotetraploid *A. arenosa* (green). Coloured bars in each sequence represent base specific SNPs relative to ancestral diploid alleles (Green = A, Blue = C, Black = G, Red = T).

## References

1. Alix K, Gerard PR, Schwarzacher T, Heslop-Harrison JSP. Polyploidy and interspecific hybridization: partners for adaptation, speciation and evolution in plants. Ann Bot. 2017;120(2):183–94. doi: 10.1093/aob/mcx079. PubMed PMID: 28854567; PubMed Central PMCID: PMC5737848.

2. Selmecki AM, Maruvka YE, Richmond PA, Guillet M, Shoresh N, Sorenson AL, et al. Polyploidy can drive rapid adaptation in yeast. Nature. 2015;519(7543):349–52. doi: 10.1038/nature14187. PubMed PMID: 25731168; PubMed Central PMCID: PMC4497379.

3. Baduel P, Bray S, Vallejo-Marin M, Kolar F, Yant L. The “Polyploid Hop”: Shifting Challenges and Opportunities Over the Evolutionary Lifespan of Genome Duplications. Front Ecol Evol. 2018;6. doi: ARTN 11710.3389/fevo.2018.00117. PubMed PMID: WOS:000451797600001.

4. Rey MD, Martin AC, Higgins J, Swarbreck D, Uauy C, Shaw P, et al. Exploiting the ZIP4 homologue within the wheat Ph1 locus has identified two lines exhibiting homoeologous crossover in wheat-wild relative hybrids. Mol Breed. 2017;37(8):95. doi: 10.1007/s11032-017-0700-2. PubMed PMID: 28781573; PubMed Central PMCID: PMC5515957.

5. Gonzalo A, Lucas MO, Charpentier C, Sandmann G, Lloyd A, Jenczewski E. Reducing MSH4 copy number prevents meiotic crossovers between non-homologous chromosomes in Brassica napus. Nat Commun. 2019;10(1):2354. doi: 10.1038/s41467-019-10010-9. PubMed PMID: 31142748; PubMed Central PMCID: PMC6541637.

6. Jenczewski E, Eber F, Grimaud A, Huet S, Lucas MO, Monod H, et al. PrBn, a major gene controlling homeologous pairing in oilseed rape (Brassica napus) haploids. Genetics. 2003;164(2):645–53. PubMed PMID: 12807785; PubMed Central PMCID: PMC1462591.

7. Henry IM, Dilkes BP, Tyagi A, Gao J, Christensen B, Comai L. The BOY NAMED SUE quantitative trait locus confers increased meiotic stability to an adapted natural allopolyploid of Arabidopsis. Plant Cell. 2014;26(1):181–94. doi: 10.1105/tpc.113.120626. PubMed PMID: 24464296; PubMed Central PMCID: PMC3963567.

8. Yant L, Hollister JD, Wright KM, Arnold BJ, Higgins JD, Franklin FCH, et al. Meiotic adaptation to genome duplication in Arabidopsis arenosa. Curr Biol. 2013;23(21):2151–6. Epub 2013/10/22. doi: 10.1016/j.cub.2013.08.059. PubMed PMID: 24139735; PubMed Central PMCID: PMCPMC3859316.

9. Page SL, Hawley RS. The genetics and molecular biology of the synaptonemal complex. Annu Rev Cell Dev Biol. 2004;20:525–58. doi: 10.1146/annurev.cellbio.19.111301.155141. PubMed PMID: 15473851.

10. Lam WS, Yang X, Makaroff CA. Characterization of Arabidopsis thaliana SMC1 and SMC3: evidence that AtSMC3 may function beyond chromosome cohesion. J Cell Sci. 2005;118(Pt 14):3037–48. doi: 10.1242/jcs.02443. PubMed PMID: 15972315.

11. Pradillo M, Knoll A, Oliver C, Varas J, Corredor E, Puchta H, et al. Involvement of the Cohesin Cofactor PDS5 (SPO76) During Meiosis and DNA Repair in Arabidopsis thaliana. Front Plant Sci. 2015;6:1034. doi: 10.3389/fpls.2015.01034. PubMed PMID: 26648949; PubMed Central PMCID: PMC4664637.

12. Bhatt AM, Lister C, Page T, Fransz P, Findlay K, Jones GH, et al. The DIF1 gene of Arabidopsis is required for meiotic chromosome segregation and belongs to the REC8/RAD21 cohesin gene family. Plant J. 1999;19(4):463–72. doi: DOI 10.1046/j.1365-313X.1999.00548.x. PubMed PMID: WOS:000082879400010.

13. Cai X, Dong F, Edelmann RE, Makaroff CA. The Arabidopsis SYN1 cohesin protein is required for sister chromatid arm cohesion and homologous chromosome pairing. J Cell Sci. 2003;116(Pt 14):2999–3007. doi: 10.1242/jcs.00601. PubMed PMID: 12783989.

14. Chelysheva L, Diallo S, Vezon D, Gendrot G, Vrielynck N, Belcram K, et al. AtREC8 and AtSCC3 are essential to the monopolar orientation of the kinetochores during meiosis. J Cell Sci. 2005;118(Pt 20):4621–32. doi: 10.1242/jcs.02583. PubMed PMID: 16176934.

15. Lambing C, Tock AJ, Topp SD, Choi K, Kuo PC, Zhao X, et al. Interacting genomic landscapes of REC8-cohesin, chromatin and meiotic recombination in Arabidopsis thaliana. Plant Cell. 2020. Epub 2020/02/07. doi: 10.1105/tpc.19.00866. PubMed PMID: 32024691.

16. Tesse S, Bourbon HM, Debuchy R, Budin K, Dubois E, Liangran Z, et al. Asy2/Mer2: an evolutionarily conserved mediator of meiotic recombination, pairing, and global chromosome compaction. Genes Dev. 2017;31(18):1880–93. Epub 2017/10/13. doi: 10.1101/gad.304543.117. PubMed PMID: 29021238; PubMed Central PMCID: PMCPMC5695089.

17. De Muyt A, Pereira L, Vezon D, Chelysheva L, Gendrot G, Chambon A, et al. A high throughput genetic screen identifies new early meiotic recombination functions in Arabidopsis thaliana. Plos Genet. 2009;5(9):e1000654. doi: 10.1371/journal.pgen.1000654. PubMed PMID: 19763177; PubMed Central PMCID: PMC2735182.

18. Armstrong SJ, Caryl AP, Jones GH, Franklin FC. Asy1, a protein required for meiotic chromosome synapsis, localizes to axis-associated chromatin in Arabidopsis and Brassica. J Cell Sci. 2002;115(Pt 18):3645–55. doi: 10.1242/jcs.00048. PubMed PMID: 12186950.

19. Ferdous M, Higgins JD, Osman K, Lambing C, Roitinger E, Mechtler K, et al. Inter-homolog crossing-over and synapsis in Arabidopsis meiosis are dependent on the chromosome axis protein AtASY3. Plos Genet. 2012;8(2):e1002507. doi: 10.1371/journal.pgen.1002507. PubMed PMID: 22319460; PubMed Central PMCID: PMC3271061.

20. Chambon A, West A, Vezon D, Horlow C, De Muyt A, Chelysheva L, et al. Identification of ASYNAPTIC4, a Component of the Meiotic Chromosome Axis. Plant Physiol. 2018;178(1):233–46. doi: 10.1104/pp.17.01725. PubMed PMID: 30002256; PubMed Central PMCID: PMC6130017.

21. Caryl AP, Armstrong SJ, Jones GH, Franklin FC. A homologue of the yeast HOP1 gene is inactivated in the Arabidopsis meiotic mutant asy1. Chromosoma. 2000;109(1-2):62–71. Epub 2000/06/16. doi: 10.1007/s004120050413. PubMed PMID: 10855496.

22. West AMV, Rosenbereg SC, Ur SN, Lehmern MK, Ye QZ, Hagemann G, et al. A conserved filamentous assembly underlies the structure of the meiotic chromosome axis. Elife. 2019;8. doi: ARTN e4037210.7554/eLife.40372. PubMed PMID: WOS:000457098200001.

23. Higgins JD, Sanchez-Moran E, Armstrong SJ, Jones GH, Franklin FC. The Arabidopsis synaptonemal complex protein ZYP1 is required for chromosome synapsis and normal fidelity of crossing over. Genes Dev. 2005;19(20):2488–500. doi: 10.1101/gad.354705. PubMed PMID: 16230536; PubMed Central PMCID: PMC1257403.

24. Schmickl R, Koch MA. Arabidopsis hybrid speciation processes. Proc Natl Acad Sci U S A. 2011;108(34):14192–7. Epub 2011/08/10. doi: 10.1073/pnas.1104212108. PubMed PMID: 21825128; PubMed Central PMCID: PMCPMC3161561.

25. Marburger S, Monnahan P, Seear PJ, Martin SH, Koch J, Paajanen P, et al. Interspecific introgression mediates adaptation to whole genome duplication. Nat Commun. 2019;10(1):5218. Epub 2019/11/20. doi: 10.1038/s41467-019-13159-5. PubMed PMID: 31740675; PubMed Central PMCID: PMCPMC6861236.

26. Hohmann N, Koch MA. An Arabidopsis introgression zone studied at high spatio-temporal resolution: interglacial and multiple genetic contact exemplified using whole nuclear and plastid genomes. BMC Genomics. 2017;18(1):810. Epub 2017/10/24. doi: 10.1186/s12864-017-4220-6. PubMed PMID: 29058582; PubMed Central PMCID: PMCPMC5651623.

27. Chelysheva L, Vezon D, Chambon A, Gendrot G, Pereira L, Lemhemdi A, et al. The Arabidopsis HEI10 is a new ZMM protein related to Zip3. Plos Genet. 2012;8(7):e1002799. Epub 2012/07/31. doi: 10.1371/journal.pgen.1002799. PubMed PMID: 22844245; PubMed Central PMCID: PMCPMC3405992.

28. Wright KM, Arnold B, Xue K, Surinova M, O’Connell J, Bomblies K. Selection on meiosis genes in diploid and tetraploid Arabidopsis arenosa. Mol Biol Evol. 2015;32(4):944–55. doi: 10.1093/molbev/msu398. PubMed PMID: 25543117; PubMed Central PMCID: PMC4379401.

29. Grishaeva TM, Bogdanov YF. Conservation and variability of synaptonemal complex proteins in phylogenesis of eukaryotes. Int J Evol Biol. 2014;2014:856230. Epub 2014/08/26. doi: 10.1155/2014/856230. PubMed PMID: 25147749; PubMed Central PMCID: PMCPMC4132317.

30. Brakenhoff RH, Schoenmakers JG, Lubsen NH. Chimeric cDNA clones: a novel PCR artifact. Nucleic Acids Res. 1991;19(8):1949. Epub 1991/04/25. doi: 10.1093/nar/19.8.1949. PubMed PMID: 2030976; PubMed Central PMCID: PMCPMC328135.

31. Chen JM, Cooper DN, Chuzhanova N, Ferec C, Patrinos GP. Gene conversion: mechanisms, evolution and human disease. Nat Rev Genet. 2007;8(10):762–75. Epub 2007/09/12. doi: 10.1038/nrg2193. PubMed PMID: 17846636.

32. Arnold B, Kim ST, Bomblies K. Single Geographic Origin of a Widespread Autotetraploid Arabidopsis arenosa Lineage Followed by Interploidy Admixture. Mol Biol Evol. 2015;32(6):1382–95. Epub 2015/04/12. doi: 10.1093/molbev/msv089. PubMed PMID: 25862142.

33. Monnahan P, Kolar F, Baduel P, Sailer C, Koch J, Horvath R, et al. Pervasive population genomic consequences of genome duplication in Arabidopsis arenosa. Nat Ecol Evol. 2019;3(3):457–68. doi: 10.1038/s41559-019-0807-4. PubMed PMID: 30804518.

34. Hazarika MH, Rees H. Genotypic control of chromosome behaviour in rye X. Chromosome pairing and fertility in autotetraploids. Heredity. 1967;22(3):317–32. doi: 10.1038/hdy.1967.44.

35. He W, Rao H, Tang S, Bhagwat N, Kulkarni DS, Ma Y, et al. Regulated Proteolysis of MutSgamma Controls Meiotic Crossing Over. Mol Cell. 2020. Epub 2020/03/05. doi: 10.1016/j.molcel.2020.02.001. PubMed PMID: 32130890.

36. Osman K, Yang J, Roitinger E, Lambing C, Heckmann S, Howell E, et al. Affinity proteomics reveals extensive phosphorylation of the Brassica chromosome axis protein ASY1 and a network of associated proteins at prophase I of meiosis. Plant J. 2018;93(1):17–33. doi: 10.1111/tpj.13752. PubMed PMID: 29078019; PubMed Central PMCID: PMC5767750.

37. Sanchez-Moran E, Santos JL, Jones GH, Franklin FC. ASY1 mediates AtDMC1-dependent interhomolog recombination during meiosis in Arabidopsis. Genes Dev. 2007;21(17):2220–33. doi: 10.1101/gad.439007. PubMed PMID: 17785529; PubMed Central PMCID: PMC1950860.

38. He Y, Wang C, Higgins JD, Yu JP, Zong J, Lu PL, et al. MEIOTIC F-BOX Is Essential for Male Meiotic DNA Double-Strand Break Repair in Rice. Plant Cell. 2016;28(8):1879–93. doi: 10.1105/tpc.16.00108. PubMed PMID: WOS:000386169200011.

39. Rao HB, Qiao H, Bhatt SK, Bailey LR, Tran HD, Bourne SL, et al. A SUMO-ubiquitin relay recruits proteasomes to chromosome axes to regulate meiotic recombination. Science. 2017;355(6323):403-7. Epub 2017/01/07. doi: 10.1126/science.aaf6407. PubMed PMID: 28059716; PubMed Central PMCID: PMCPMC5569317.

40. Ahuja JS, Sandhu R, Mainpal R, Lawson C, Henley H, Hunt PA, et al. Control of meiotic pairing and recombination by chromosomally tethered 26S proteasome. Science. 2017;355(6323):408-11. Epub 2017/01/07. doi: 10.1126/science.aaf4778. PubMed PMID: 28059715; PubMed Central PMCID: PMCPMC6054871.

41. Zhang L, Wang S, Yin S, Hong S, Kim KP, Kleckner N. Topoisomerase II mediates meiotic crossover interference. Nature. 2014;511(7511):551-6. doi: 10.1038/nature13442. PubMed PMID: 25043020; PubMed Central PMCID: PMC4128387.

42. Jorgensen MH, Ehrich D, Schmickl R, Koch MA, Brysting AK. Interspecific and interploidal gene flow in Central European Arabidopsis (Brassicaceae). BMC Evol Biol. 2011;11:346. doi: 10.1186/1471-2148-11-346. PubMed PMID: 22126410; PubMed Central PMCID: PMC3247304.

43. Bomblies K, Higgins JD, Yant L. Meiosis evolves: adaptation to external and internal environments. The New phytologist. 2015;208(2):306–23. doi: 10.1111/nph.13499. PubMed PMID: 26075313.

44. Giraut L, Falque M, Drouaud J, Pereira L, Martin OC, Mezard C. Genome-Wide Crossover Distribution in Arabidopsis thaliana Meiosis Reveals Sex-Specific Patterns along Chromosomes. Plos Genet. 2011;7(11). doi: ARTN e100235410.1371/journal.pgen.1002354. PubMed PMID: WOS:000297264500009.

45. Otto SP. The evolutionary consequences of polyploidy. Cell. 2007;131(3):452–62. doi: 10.1016/j.cell.2007.10.022. PubMed PMID: 17981114.

46. Hu TT, Pattyn P, Bakker EG, Cao J, Cheng JF, Clark RM, et al. The Arabidopsis lyrata genome sequence and the basis of rapid genome size change. Nat Genet. 2011;43(5):476-+. doi: 10.1038/ng.807. PubMed PMID: WOS:000289972600020.

47. Edgar RC. MUSCLE: multiple sequence alignment with high accuracy and high throughput. Nucleic Acids Res. 2004;32(5):1792–7. doi: 10.1093/nar/gkh340. PubMed PMID: 15034147; PubMed Central PMCID: PMC390337.

48. Mascher M, Gundlach H, Himmelbach A, Beier S, Twardziok SO, Wicker T, et al. A chromosome conformation capture ordered sequence of the barley genome. Nature. 2017;544(7651):427-33. doi: 10.1038/nature22043. PubMed PMID: 28447635.

49. Higgins JD, Wright KM, Bomblies K, Franklin FC. Cytological techniques to analyze meiosis in Arabidopsis arenosa for investigating adaptation to polyploidy. Front Plant Sci. 2014;4:546. doi: 10.3389/fpls.2013.00546. PubMed PMID: 24427164; PubMed Central PMCID: PMC3879461.

50. Chelysheva L, Grandont L, Vrielynck N, le Guin S, Mercier R, Grelon M. An easy protocol for studying chromatin and recombination protein dynamics during Arabidopsis thaliana meiosis: immunodetection of cohesins, histones and ML H1. Cytogenet Genome Res. 2010;129(1-3):143-53. Epub 2010/07/16. doi: 10.1159/000314096. PubMed PMID: 20628250.

51. Lambing C, Osman K, Nuntasoontorn K, West A, Higgins JD, Copenhaver GP, et al. Arabidopsis PCH2 Mediates Meiotic Chromosome Remodeling and Maturation of Crossovers. Plos Genet. 2015;11(7):e1005372. doi: 10.1371/journal.pgen.1005372. PubMed PMID: 26182244; PubMed Central PMCID: PMC4504720.

52. Wong YH, Lee TY, Liang HK, Huang CM, Wang TY, Yang YH, et al. KinasePhos 2.0: a web server for identifying protein kinase-specific phosphorylation sites based on sequences and coupling patterns. Nucleic Acids Research. 2007;35:W588–W94. doi: 10.1093/nar/gkm322. PubMed PMID: WOS:000255311500110.

53. Blom N, Sicheritz-Ponten T, Gupta R, Gammeltoft S, Brunak S. Prediction of post-translational glycosylation and phosphorylation of proteins from the amino acid sequence. Proteomics. 2004;4(6):1633–49. doi: 10.1002/pmic.200300771. PubMed PMID: 15174133.

54. Zhao Q, Xie YB, Zheng YY, Jiang S, Liu WZ, Mu WP, et al. GPS-SUMO: a tool for the prediction of sumoylation sites and SUMO-interaction motifs. Nucleic Acids Research. 2014;42(W1):W325–W30. doi: 10.1093/nar/gku383. PubMed PMID: WOS:000339715000054.

55. Kahle D,Wickham H. ggmap: Spatial Visualization with ggplot2. R J. 2013;5(1):144–61. PubMed PMID: WOS:000321944400015.

